# Phenotypic variation and quantitative trait loci for resistance to southern anthracnose and clover rot in red clover

**DOI:** 10.1101/2022.05.23.493028

**Authors:** Lea A. Frey, Tim Vleugels, Tom Ruttink, Franz X. Schubiger, Marie Pegard, Leif Skøt, Christoph Grieder, Bruno Studer, Isabel Roldán-Ruiz, Roland Kölliker

## Abstract

Red clover (*Trifolium pratense* L.) is an important forage legume of temperate regions, particularly valued for its high yield potential and its high forage quality. Despite substantial breeding progress during the last decades, continuous improvement of cultivars is crucial to ensure yield stability in view of newly emerging diseases or changing climatic conditions. The high amount of genetic diversity present in red clover ecotypes, landraces and cultivars provides an invaluable, but often unexploited resource for the improvement of key traits such as yield, quality, and resistance to biotic and abiotic stresses.

A collection of 397 red clover accessions was genotyped using a pooled genotyping-by-sequencing approach with 200 plants per accession. Resistance to the two most pertinent diseases in red clover production, southern anthracnose caused by *Colletotrichum trifolii*, and clover rot caused by *Sclerotinia trifoliorum*, was assessed using spray inoculation. The mean survival rate for southern anthracnose was 22.9% and the mean resistance index for clover rot was 34.0%. Genome-wide association analysis revealed several loci significantly associated with resistance to southern anthracnose and clover rot. Most of these loci are in coding regions. One quantitative trait locus (QTL) on chromosome 1 explained 16.8% of the variation in resistance to southern anthracnose. For clover rot resistance we found eight QTL, explaining together 80.2% of the total phenotypic variation. The SNPs associated with these QTL provide, once validated, a promising resource for marker-assisted selection in existing breeding programs, facilitating the development of novel cultivars with increased resistance against two devastating fungal diseases of red clover.

**Key message:** High variability for and candidate loci associated with resistance to southern anthracnose and clover rot in a worldwide collection of red clover provide a first basis for genomics-assisted breeding.

## Introduction

Red clover (*Trifolium pratense* L.), one of the most important forage legumes in temperate climates, is grown in mixture with forage species or as a pure stand (Taylor 2008). Red clover is appreciated for its high forage yield (up to 14 tons dry matter/ha/year), its high protein content and digestibility, its ability to fix atmospheric nitrogen, and its beneficial effects on soil structure (Broderick 1995; Taylor and Quesenberry 1996; Halling et al. 2004; Nyfeler et al. 2011). Red clover is an outcrossing species with a high degree of self-incompatibility and a genome size of approximately 420 Mb (2n = 2x = 14; De Vega et al. 2015). Main breeding objectives are a high and stable forage yield, persistence, and good forage quality. Disease and insect resistance are important aspects of red clover breeding programs, and necessary to meet the requirements of a successful cultivar (Taylor 2008; Boller et al. 2010).

Key fungal pathogens threatening European red clover production and leading to severe yield losses are *Colletotrichum trifolii* Bain & Essary, causing southern anthracnose, and *Sclerotinia trifoliorum* Erikks, causing clover rot. *C. trifolii* was first described in 1906 and has since been reported on a regular basis in most European countries (Bain and Essary 1906; Schubiger et al. 2004; Jacob et al. 2015). In the southern parts of the US, where the disease has long been a major problem, intensive breeding efforts led to largely resistant cultivars (Taylor 2008). *C. trifolii* has benefitted from warmer summer temperatures in Central Europe, and southern anthracnose has become a limiting factor for red clover production, increasing the demand for resistant cultivars (Boller et al. 2010). *C. trifolii* is a hemibiotrophic fungus that mainly spreads by rain and wind and causes brown coloration on petioles and stems of red clover. Once the xylem is infected the plant begins to shrivel, stem lesions occur, and the plant eventually dies off (De Silva et al. 2017). While the genetics of southern anthracnose resistance in red clover remains largely unknown, resistance to *Colletotrichum* spp. has been extensively studied in other plant species, including soybean (*Glycine max* L.), common bean (*Phaseolus vulgaris* L.), and alfalfa (*Medicago sativa* L.), as reviewed in Dean et al. (2012).

Clover rot also known as *Sclerotinia* crown, stem rot or clover cancer is caused by the necrotrophic fungus *S. trifoliorum*, which can survive up to seven years as soil-borne resting bodies (sclerotia). In autumn, sclerotia develop apothecia, which release airborne ascospores that infect red clover leaves and slowly colonize the whole plant during winter (Taylor and Quesenberry 1996; Öhberg 2008). Prolonged conditions of high humidity such as temperate, damp weather or long periods of snow cover favor clover rot development (Saharan and Mehta 2010). Although little is known on its genetics, resistance to clover rot in red clover is assumed to be a quantitative trait (Poland et al. 2009; Klimenko et al. 2010; Vleugels and van Bockstaele 2013).

As southern anthracnose and clover rot can cause substantial losses in European red clover production, resistance breeding is of prime importance. Different aspects need to be considered in developing resistant cultivars. First, as for most diseases, natural infection typically varies between years and between locations. Therefore, infection in breeding trials is rarely homogeneous, and disease development strongly depends on weather conditions. Second, most red clover cultivars are bred as synthetic, population-based varieties, complicating the fixation of resistance alleles. Third, little is known on the genetic basis of resistance against southern anthracnose or clover rot, precluding the use of molecular markers in resistance breeding. Resistance breeding for southern anthracnose, and for clover rot to a lesser extent, has been relatively successful when using artificial inoculations or bio-tests in controlled environments (Marum et al. 1994; Delclos and Duc 1996; Schubiger et al. 2003, 2004; Vleugels and van Bockstaele 2013; Hartmann et al. 2022). However, DNA markers reliably predicting resistance to both diseases would allow to substantially save time, effort and resources through genomic prediction and early generation marker-assisted selection (MAS) (Collard and Mackill 2008). Furthermore, MAS allows to combine multiple favored alleles through fewer crossing events when compared to pure phenotypic selection (Collard and Mackill 2008).

The main objective of this study was to better characterize disease resistance for the two most relevant fungal diseases threatening red clover production in Europe and other temperate zones worldwide and to identify genetic loci linked to resistance. Therefore, we screened a diverse collection of red clover accessions under controlled conditions for southern anthracnose and clover rot resistance. We examined the phenotypic variation in resistance to these two diseases, aiming to find accessions with a high degree of resistance to one or both diseases. Furthermore, we developed genome-wide allele frequency fingerprints using pooled genotyping-by sequencing (pool-GBS) and performed genome-wide association studies (GWAS) to identify quantitative trait loci (QTL). Potential candidate resistance genes were identified in the genomic regions underlying the QTL associated with southern anthracnose and clover rot resistance.

## Material and Methods

We used a collection of 397 red clover accessions that was established in the frame of the EUCLEG project (Horizon 2020 Programme for Research & Innovation, grant agreement n°727312; http://www.eucleg.eu). This collection (hereafter referred to as the EUCLEG-accessions) contains plant material from 23 countries including cultivars, breeding material, landraces, and ecotypes. Detailed information on the EUCLEG-accessions is given in Supplementary Table S1. Each EUCLEG-accession can be considered as a population of related plants. While all accessions were used for genotyping and phenotyping of southern anthracnose resistance, only 392 accessions were screened for clover rot resistance.

### Genotyping and filtering for single nucleotide polymorphisms (SNPs)

Seedlings were grown in the greenhouse in 96-compartment plant trays filled with compost. At the one-leaf stage, that leaf was harvested from 200 seedlings per accession. Fresh leaves from the same accession were pooled and DNA was extracted using the QIAGEN DNeasy 96 Plant kit (QIAGEN, Citylabs 2.0, Manchester M13 0BH, UK). The DNA concentration was measured using a Qubit(tm)2.0 instrument and normalized to 20 ng µl^-1^. Genotyping was realized by LGC Genomics (Berlin, Germany) using a *Pst*I-*Mse*I double digest pool-GBS method, in combination with PE-150 Illumina sequencing. SNP calling and allele frequency calculations were done as described in Keep et al. (2020). A detailed description of the parameters specific for this study is provided in the Supplementary Methods. Data were filtered to retain SNPs with less than 5% missing values, allele frequencies between 0.05 and 0.95 in at least 10 accessions and mean allele frequencies across all accessions between 0.05 and 0.95 (0.05 < MAF < 0.95). Missing data was replaced by the mean allele frequency across all accessions per SNP. The GBS reads are available in NCBI SRA.

### Experimental design and phenotyping

#### Southern anthracnose

A resolvable row-column design with two standard cultivars as controls (‘Pavo’ and ‘Milvus’), and four full replications was used. Experimental units consisted of 24 plants of the same accession sown together. Plants were grown in plastic boxes (300 × 400 × 145 mm) filled with cultivation substrate at a plant-to-plant distance of approx. 4 cm. One replicate consisted of 160 boxes, each box containing 72 plants of three different accessions (3 × 24 = 72).

Spray-inoculation was adapted from Schubiger et al. (2003). Briefly, plants were grown in a greenhouse (19-23°C, 16 h light from sodium-vapor bulbs, >100 μEm^−2^ s^−1^) at Agroscope (Zurich, Switzerland). After six weeks, plants were cut 3-4 cm above the ground and allowed to regrow for two weeks. The number of living plants per experimental unit (G) was determined before plants were inoculated with a single-spore isolate. Fungal spores of the isolate CTR 010103 (collected on red clover in 2001 in Ellighausen, Switzerland) were grown on potato dextrose agar (PDA) at around 18°C in the dark and 12 h ultraviolet light per 24 h. After ten days, spores were gently removed with sterile dH_2_O. The concentration of the spore suspension was adjusted with dH_2_O to 3.2-4.8 × 10^6^ spores ml^-1^ by counting spores under the microscope. For inoculation, approximately 40 ml spore suspension was used per box, wetting the plants from top to bottom using a spray gun compressor at 2 bar. The inoculated plants were covered with a polyethylene sheet for five days. Plants were cut four times at 14-, 42-, 70- and 98-days post inoculation (dpi). The survival rate (S_rate_) was assessed by counting the surviving plants (S) two weeks after the second cut (56 dpi) multiplied by 100 and divided by the total number of plants before inoculation (G, Equation 1).

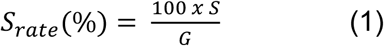

After scoring, the surviving plants were re-inoculated with a mixture of seven additional *C. trifolii* single-spore isolates collected 2001 in Ellighausen (CTR010101, CTR010102, CTR010104, CTR010105, CTR010106, CTR010107, CTR010108). Inoculation was conducted as described above.

Seven weeks after the second inoculation (105 days after the first inoculation), the surviving plants were counted. Cumulative survival rate (CumS_rate_) was calculated as survivors after the second inoculation period (S_cum_) divided by G (Equation 2).

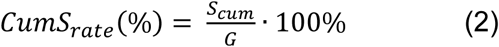

#### Clover rot

A total of 13 separate clover rot trials were performed, so that all accessions were screened in three full replicates. These trials comprised up to 52 trays containing 94 accessions with 36 plants each, along with two positive and two negative (non-inoculated) control trays (both fully sown with the control cultivar ‘Lemmon’). Plants were sown eight weeks prior to inoculation in Quickpot^®^ trays (HerkuPlast QP96T, InterGrow, Alter, Belgium) in peat substrate (Saniflor Beroepspotgrond, InterGrow, Aalter, Belgium). Each tray was seeded with three accessions: one in the three top rows (36 plants), one in the three bottom rows (36 plants), and the control cultivar ‘Lemmon’ in the two middle rows (24 plants). Plants were grown in the greenhouse (20-25°C, 12 h light from TL lamps at 50 μEm^−2^ s^−1^) and watered when required. Three weeks prior to inoculation, plants were cut at 5 cm above the ground. A single-spore isolate derived from Cz.A 1 was chosen for further experiments, as it possessed the highest growth speed on PDA medium (Vleugels et al. 2013a). Inoculum was prepared to contain approximately 8,000 mycelium fragments ml^-1^ in sterile dH_2_O with 5 g l^-1^ glucose and 150 µl l^-1^ Tween 20 (Sigma-Aldrich, Germany). Two to four days prior to inoculation, trays were moved to a growth chamber (15°C, 12 h light), where they were randomly placed on eight growing tables and watered until saturation. After inoculation, the tables were covered with caps made of transparent plastic foil and misted to increase humidity. Plants were sprayed with mycelium suspension until run-off, after which the plastic caps were closed, and the lights dimmed until the next morning. The negative control trays were sprayed with infection solution without inoculum. Water was misted over the plants at day three and day six dpi and the plastic foil was replaced immediately after misting. After nine days of incubation, the plastic foil was removed, and the disease incidence was scored on each plant using a scale from 1 (no symptoms) to 5 (completely dead plant). Subsequently, scores were converted into percentages through calculation of the resistance index (RI) adapted from Marum et al. (1994) as follows (Equation 3).

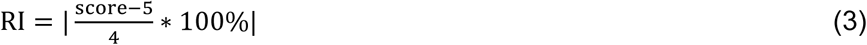

### Calculation of mean values per accession and heritabilities

Statistical analyses were carried out in R statistical software version 4.0.3 (R Core Team 2021) and RStudio version 1.3.1093 (RStudio Team 2020) and the mixed model package ASReml-R version 4.0 (Butler et al. 2017). Assumptions of homoscedasticity of variances and normality of residuals were met according to residual plots, except for the cumulative survival rate, which was thus square root transformed. Linear mixed model analyses for southern anthracnose resistance were performed using accessions and replicates as fixed effects and all other parameters as random effects in the model (Equation 4).

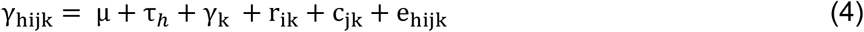

where γ_hijk_ is the survival rate or cumulative survival rate of the *h*-th accession in the *i*-th row and *j*-th column nested within *k*-th complete replicate, μ is the general mean, τ_h_ the effect of the *h*-th accession, γ_k_ the effect of *k*-th complete replicate, r_ik_ the effect of *i*-th row within *k*-th replicate, c_jk_ the effect of *j*-th column within *k*-th replicate, and e_hijk_ the residual error per experimental unit.

Linear mixed model analysis for clover rot resistance was performed using accessions as fixed effects and all other parameters as random effects in the model (Equation 5).

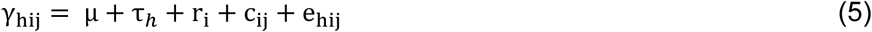

where γ_hij_ is the resistance index of the *h*-th accession on the *j*-th table nested within the *i-*th trial, μ is the general mean, τ_h_ the effect of the *h*-th accession, r_i_ the effect of *i*-th trial, c_ij_ the effect of *j*-th table within *i*-th trial, and e_hij_ the residual error per experimental unit.

For fixed effects Wald *x*^*2*^ tests with ssType = “conditional” were performed. Best linear unbiased estimates (BLUEs) for each accession and all traits were calculated. Heritability was calculated according to Cullis et al. (2006, Equation 6).

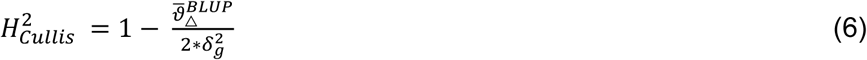

where d^2^ is the variance of the accession and 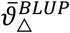 the average standard error of the accession BLUPs. Pairwise Wilcoxon rank sum tests were performed with a Bonferroni threshold of α = 5% to compare the different red clover accession types (i.e., ecotypes, landraces, breeding material, and cultivars). BLUEs were used as accession values for downstream analyses.

### Genomic relationship matrix and association between SNPs and phenotypic traits

The genomic relationship matrix was calculated as described in Cericola et al. (2018) with a ploidy number of 16, which is assumed to be ideal when dealing with synthetic cultivars. GWAS were performed with the multi-locus mixed-model (MLMM) approach implemented in the R package mlmm.gwas (Segura et al. 2012; Bonnafous et al. 2019). Through forward inclusion and backward elimination, SNPs were integrated as cofactors into a mixed-model regression approach. Variance components of the model were estimated at each step separately. The number of steps were limited to 20, and the model with the lowest Bayesian information criterion (BIC) was selected (Chen and Chen 2008). The percentage of phenotypic variation explained by each SNP was obtained by comparing the R2 of a linear model taking SNPs as fixed effects and the kinship matrix as random effect to the R^2^ of the same model without integrating the SNPs. Mixed linear models were calculated with the “lmekin” function of the “coxme” R package (Therneau 2020). SNP positions and adjacent regions of the genome annotation of the red clover reference genome sequence v2.1 (De Vega et al. 2015) was visualized using CLC genomic workbench version 9 (CLC bio, Aarhus, Denmark). Sequences of the genes containing the significant SNPs (Table 2) and genes in adjacent regions (10 kb up- and downstream) were compared to the *M. truncatula* genome (BLASTn; Tang et al. 2014) and the BLAST hit with the lowest the e-value was selected.

## Results

Phenotypic variation among the 397 EUCLEG-accessions was high for southern anthracnose resistance. For the single-spore inoculation, the survival rate ranged from 0% to 79.9% (Fig.1a), and for the mixed-spore inoculation the back-transformed cumulative survival rate (square root transformed) ranged from 0% to 73.5% (Fig.1b). The overall mean was 22.9% for the single-spore inoculation and 13.1% for the mixed-spore inoculation. Mean survival rate of the rather susceptible cultivar ‘Milvus’ was 20.6% for the single-spore inoculation and 8.7% for the mixed-spore inoculation, which was only 2.3% and 4.4% lower than the overall mean of the trial for single-spore and mixed-spore inoculation, respectively (Fig. 1a, 1b). Mean survival rate of the rather resistant cultivar ‘Pavo’ (43.7% for single-spore and 29.9% for mixed-spore inoculation) was 20.8% and 16.8% higher than the mean of all accessions for single-spore and mixed-spore inoculation, respectively.

**Fig. 1.**
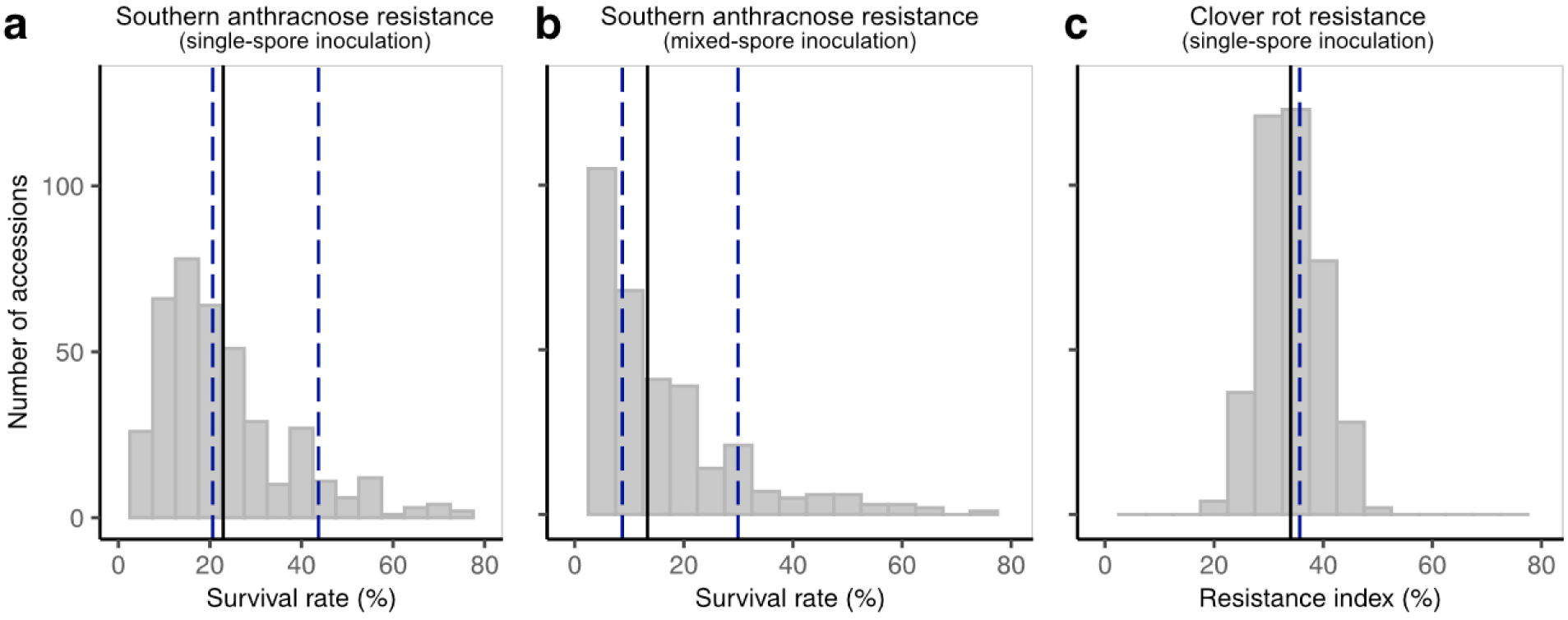
Frequency distribution of resistance scores of the red clover (*Trifolium pratense* L.) EUCLEG-accessions. Plots depict adjusted means for southern anthracnose survival rate (a), back-transformed cumulative survival rate (b), and resistance index for clover rot (c). Means are indicated by the solid line and compared with the control cultivars (dashed lines) ‘Milvus’ (left) and ‘Pavo’ (right) for southern anthracnose resistance (a and b), and ‘Lemmon’ for clover rot resistance (c).

For clover rot resistance, the phenotypic variation among the EUCLEG-accessions was smaller, with resistance indices ranging from 19.7% to 48.9% and a mean of 34.0% (Fig. 1c). The cultivar ‘Lemmon’ showed, with a resistance index of 35.7%, a similar resistance index as the average of all accessions.

Means and standard errors for the three traits and for all accessions are listed in Supplementary Table S2. Variance components for the three traits were significant for accession effects (Table 1). Heritability was moderately high, with 0.85 for southern anthracnose resistance (single-spore and mixed-spore inoculation), and 0.88 for clover rot resistance. Comparable values for heritability were obtained with the method of Piepho and Möhring (2007, data not shown). Grouping the accessions into breeding material, cultivars, landraces, and ecotypes revealed significant differences (*p* < 0.05) among the different groups (Fig. 2). For southern anthracnose, landraces showed a significantly lower survival rate compared to the other three types of material with a high variation within the four groups for single-spore inoculation (Fig. 2a). The results for the mixed-spore inoculation were comparable (data not shown). For clover rot resistance, breeding material performed slightly, but significantly (*p* < 0.05), better than cultivars, landraces, and ecotypes.

**Table 1.**
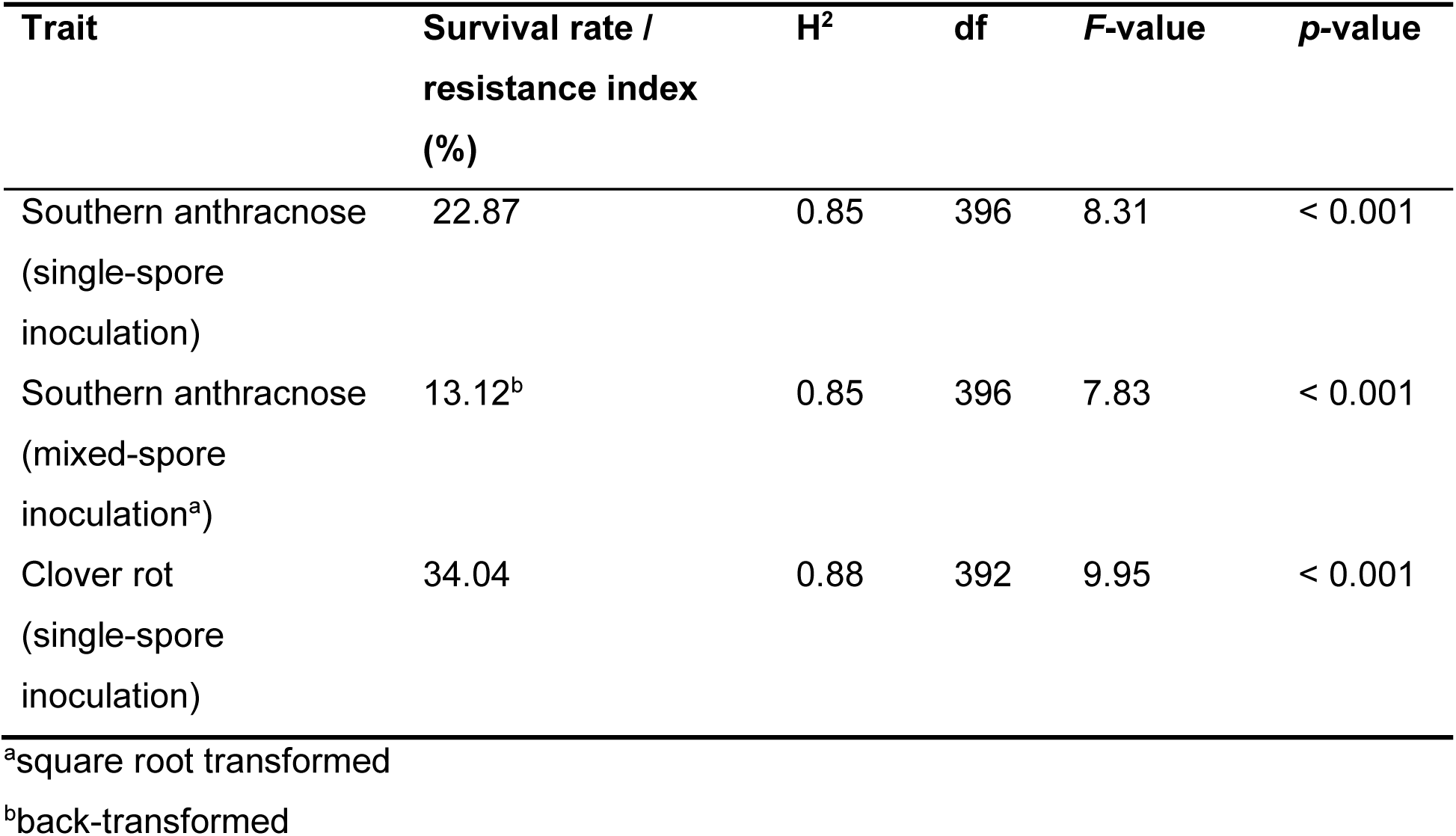
Mean survival rates for southern anthracnose, and resistance index for clover rot as well as Cullis heritabilities (H2) were performed.

**Fig. 2.**
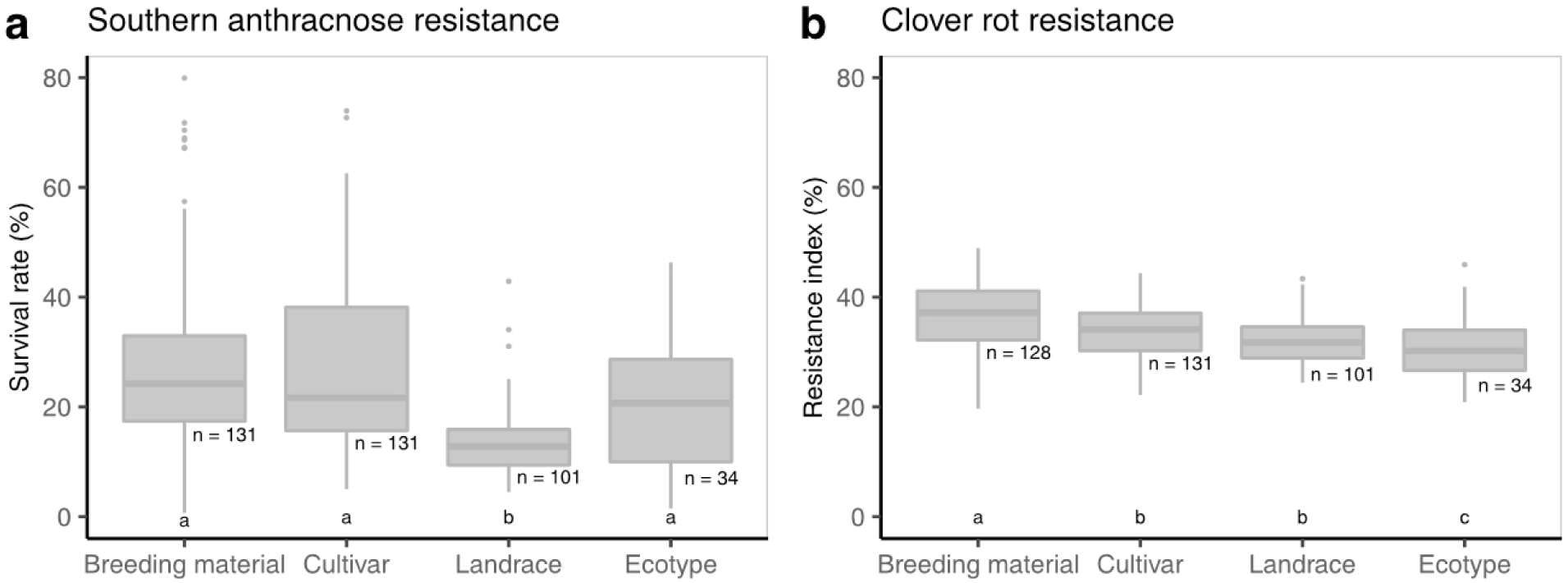
Accession means grouped according to the type of accession for survival rate after southern anthracnose single-spore inoculation (a) and resistance index for clover rot (b). Horizontal bars represent medians. Medians with no letter in common were significantly different (Kruskal-Wallis; α = 5%).

The variation in clover rot resistance within the four groups was low compared to southern anthracnose resistance and reflected the overall lower level of variation. Landraces and cultivars from the US showed a generally high resistance to southern anthracnose with the lowest survival rate being as high as 50.8%. Substantial resistance was also observed in breeding material and cultivars from Argentina, Czech Republic, and Switzerland (Fig. 3a). For clover rot resistance, breeding material from Sweden and Norway performed best with resistance indices of 48.9% and 44.6%, respectively (Fig. 3b).

**Fig. 3.**
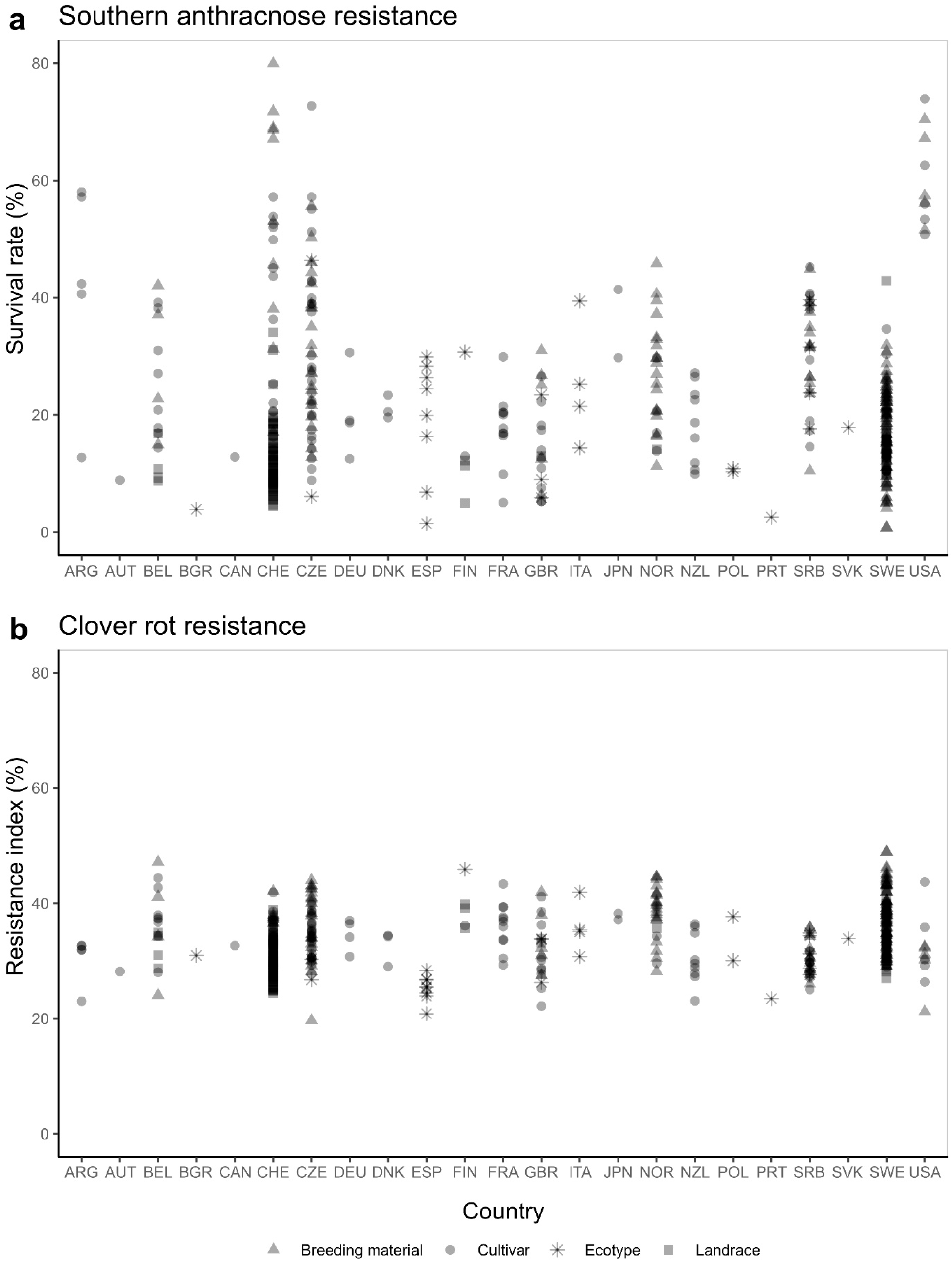
Accession means grouped according to their origin (three letter country code). Survival rates for single-spore inoculation (a) and resistance index for clover rot (b).

Genome-wide allele frequency fingerprints retained a total of 20,137 SNPs with 0.7% missing values after filtering. SNP reference allele frequencies across all accessions were biased towards one, indicating that many low-frequency alternative alleles exist while the most abundant allele across the accessions is consistent with the nucleotide encoded in the reference genome sequence (left skewed, Supplementary Fig. S1). A total of 7,372 SNPs were located on scaffolds with unassigned chromosomal position. The other 12,765 SNPs (63.4%) were evenly spread across the seven chromosomes. The average SNP density of the SNPs assigned to the seven chromosomes was 26.85 SNPs per 250 kb (Supplementary Fig. S2). Linkage disequilibrium (LD) of adjacent SNPs (Supplementary Fig. S3) was almost absent and therefore flanking genes were only identified in the 10 kb regions up- and downstream of significant SNPs.

We found several SNPs that were significantly associated with each of the three traits investigated (Fig. 4). The SNP “LG1_6601280” explained 16.8% and 14.3% of the total phenotypic variation for southern anthracnose resistance after single-spore inoculation and mixed-spore inoculation, respectively. For clover rot resistance the most relevant SNP “Scaf658_16191” explained 12.8% of the total phenotypic variation. Another five SNPs, each explaining about 10% of the phenotypic variation for clover rot resistance, were two located on chromosome 1. For all traits combined, a total of 22 SNPs were significantly associated after Bonferroni correction (α = 5%), of which 18 were found in coding regions of the reference genome. The function of these genes in red clover was assigned based on orthology to *M. truncatula* genes (Table 2).

**Fig. 4.**
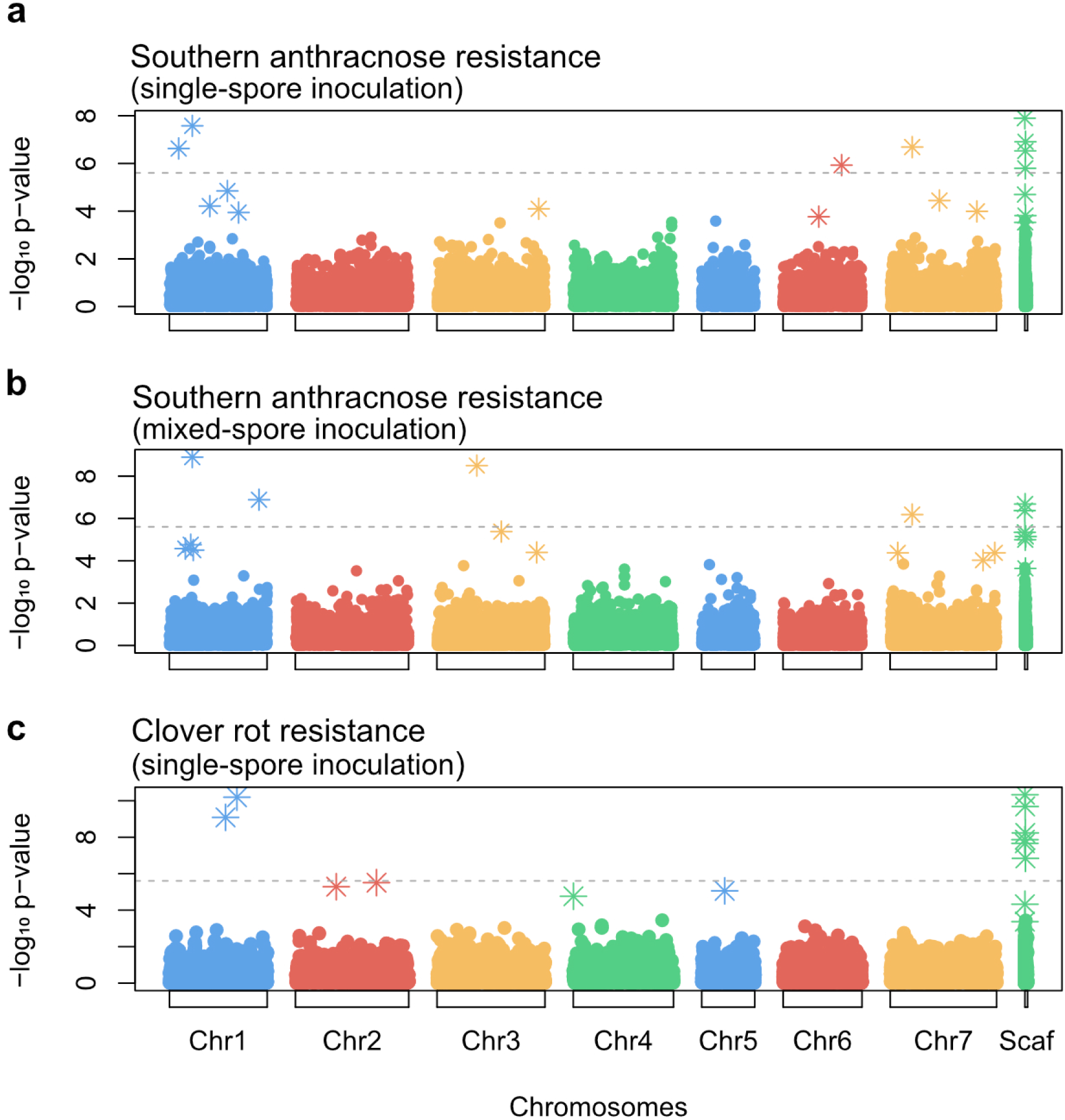
Genome-wide association study based on 20,137 single nucleotide polymorphismns using the multi-locus mixed-model approach (Segura et al., 2012), for survival rate after southern anthracnose single-spore inoculation (a), mixed-spore inoculation (b) and resistance index for clover rot (c). The dotted line represents the significance threshold after Bonferroni correction (α = 5%).

**Table 2.**
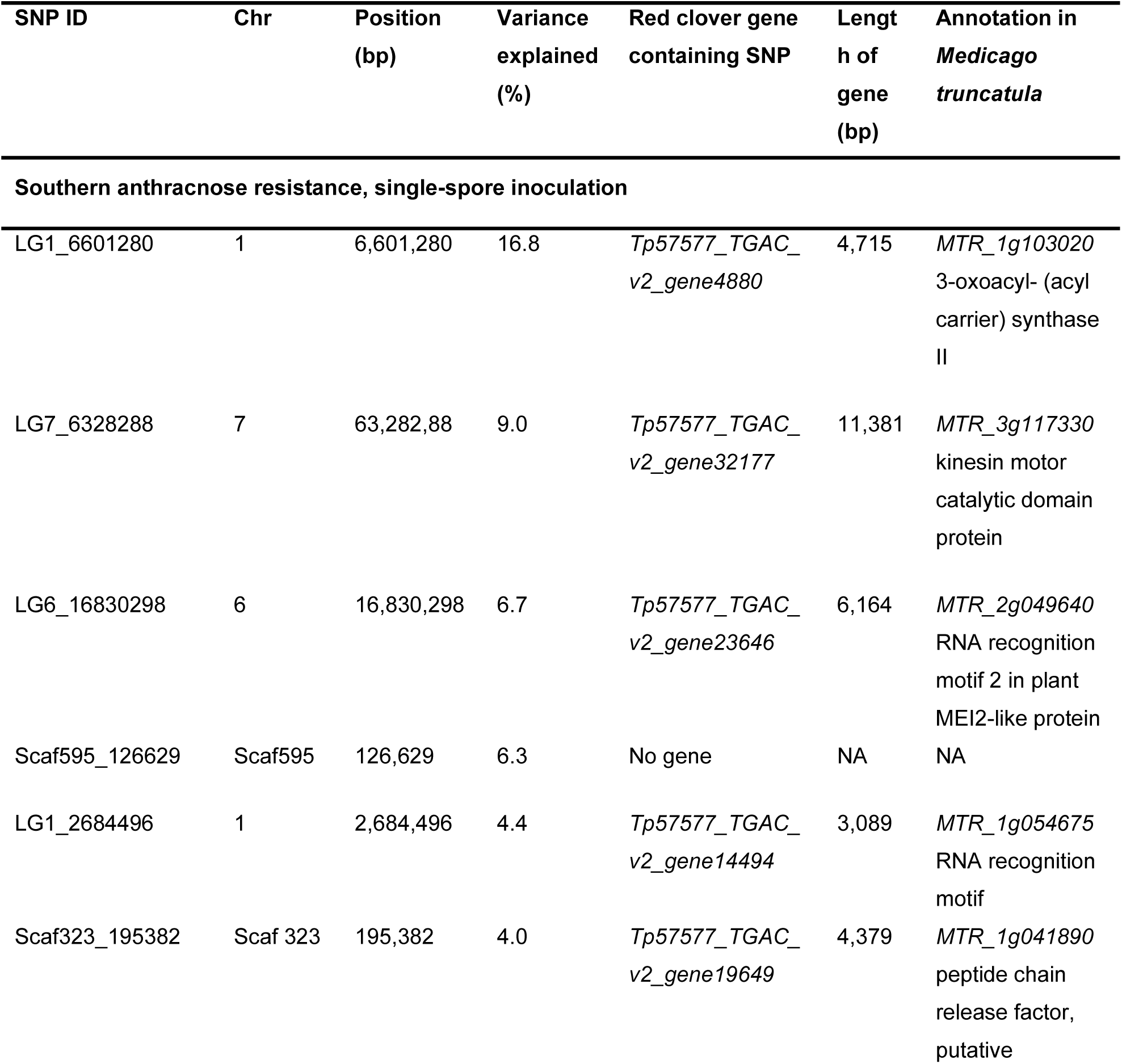
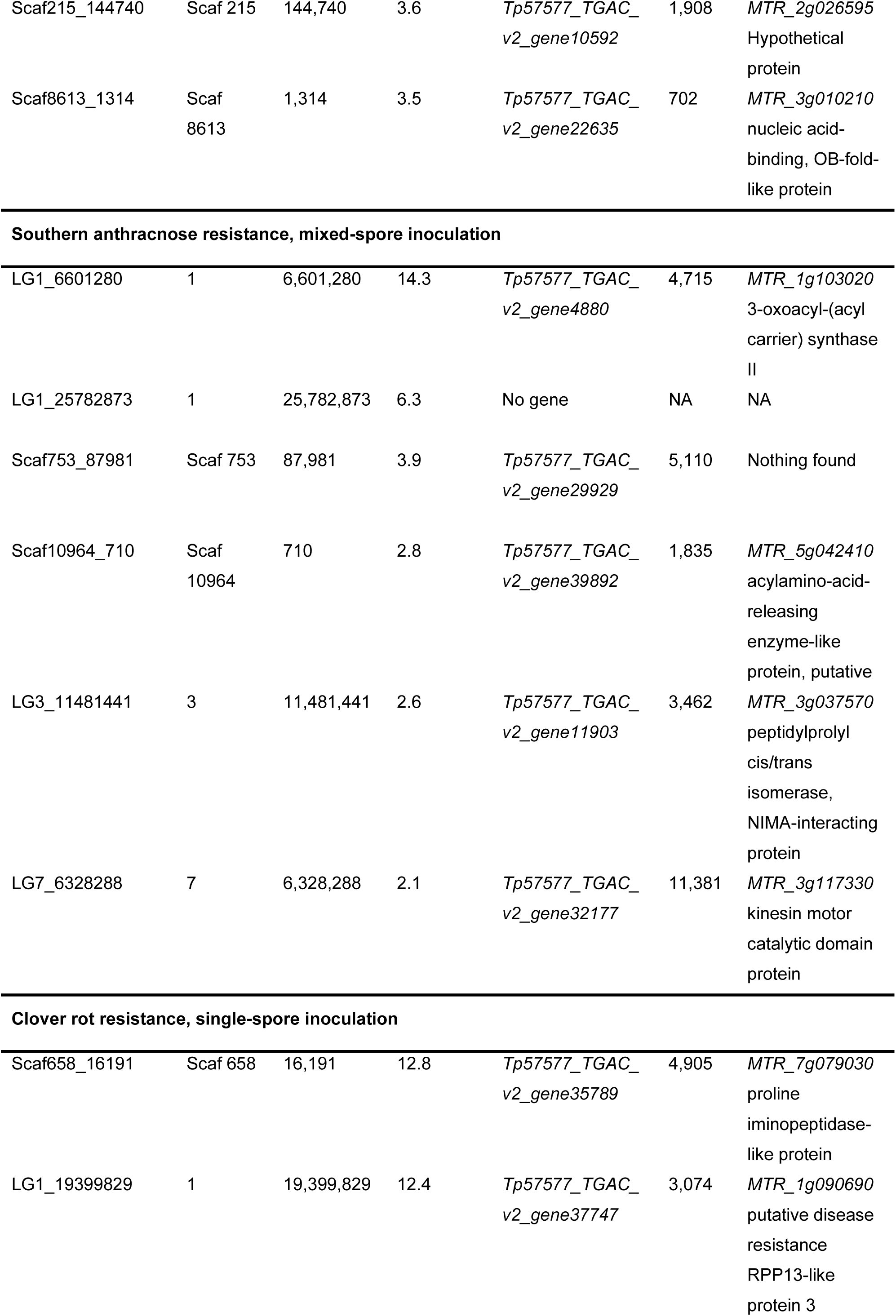
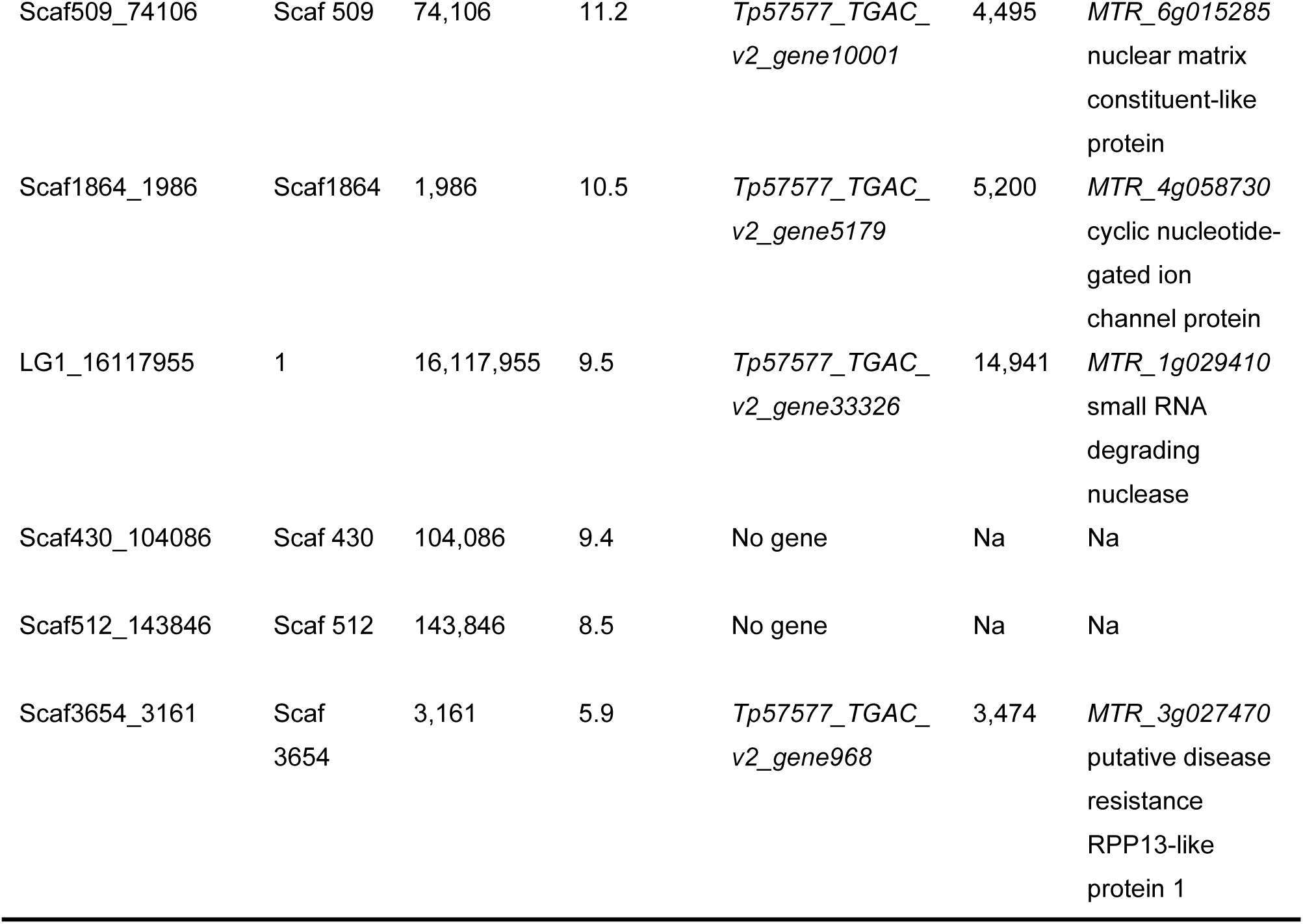
Single-nucleotide polymorphisms associated with southern anthracnose and clover rot resistance. SNP ID, chromosome (Chr), SNP position on the chromosome, phenotypic variance explained by that SNP, red clover gene containing the SNP, length in (bp) of the red clover genes, and the gene name and gene function of the closest ortholog in *M. truncatula* are listed.

## Discussion

Most of the 397 red clover accessions showed a high susceptibility to southern anthracnose and clover rot. In recent decades, summers in Central Europe became warmer and *C. trifolii* infections increased (Boller et al. 2010). Winters became damper and long periods of dry frost became less likely, conditions which favor clover rot infections (Öhberg 2008). Southern anthracnose and clover rot became a limiting factor for red clover production in regions that were previously not affected (Boller et al. 2010; Jacob et al. 2015). Changing climatic conditions can substantially shape crop pathogen assemblages (Chaloner et al. 2021). The lack of adaptation to the newly emerging pathogens may explain the overall high susceptibility for southern anthracnose and clover rot observed in the red clover EUCLEG-accessions.

In general, ecotypes from Southern European countries showed a higher survival rate after inoculation with *C. trifolii* (Fig. 3a), compared to populations from Northern latitudes. Also, US cultivars and breeding materials performed well with a high survival rate after spray-inoculation with *C. trifolii*. This may be explained by the intense selection efforts for southern anthracnose resistance in the US since the 1950s (Taylor 2008). Landraces, ecotypes, cultivars and breeding material from Sweden and Finland showed a higher resistance index for clover rot compared to populations from the other regions (Fig. 3). In Sweden, clover rot is the major cause of red clover stand failure since decades and early generation selection for clover rot resistance is indispensable (Lundin and Jönsson 1974). Some accessions from Belgium, Switzerland and the Czech Republic, mainly new breeding material, showed an increased resistance to either one or both diseases (Fig. 3), reflecting more recent attempts to actively select for southern anthracnose and clover rot resistance. These observations are consistent with the hypothesis that resistance levels increase in regions where pathogens occur, by adaptation through natural or artificial selection over time (Huxley 1939; Burdon and Thrall 2009). Both types of selection have probably played a role in shaping the geographical differentiation in levels of red clover disease resistance against the pathogens investigated in this study.

Despite the generally high susceptibility of the EUCLEG-accessions, there was considerable phenotypic variation for resistance. We observed a high variation in resistance to southern anthracnose, whereas the phenotypic variation for clover rot resistance was comparably low. For southern anthracnose, several accessions did show a high survival rate and could present a valuable resistance source for breeding programs. On the other hand, no accession showed an appropriate level for clover rot resistance, thus hampering direct introgression of resistance into existing breeding material. However, recurrent selection after artificial inoculation has previously been shown to improve levels of resistance against clover rot in red clover and resistance to *S. sclerotiorum* in different legume species (Terán and Singh 2009; Vleugels et al. 2013b). Recurrent selection after artificial inoculation trials seems to date the only option to substantially increase resistance levels for clover rot as well as for southern anthracnose in red clover (Schubiger et al. 2003; Vleugels et al. 2013b; Jacob et al. 2015).

Despite considerable success, phenotypic recurrent selection using artificial spray-inoculation is time and labor intensive and requires large greenhouse trials. Furthermore, the fixation of resistance alleles in population-based cultivars is difficult (Patella et al. 2019). DNA markers that are linked to specific QTL harboring genes with a specific role in resistance, so-called diagnostic markers, are routinely used in cultivar development of major crops like wheat (*Triticum aestivum* L.) and maize (*Zea Mays* L. ; Miedaner and Flath 2007; Guo et al. 2019). MAS is particularly effective for qualitative resistance traits where only a few genes underlying the resistance are involved (Adam-Blondon et al. 1994; Xiao et al. 2001; Zhou et al. 2001).

Current breeding programs would benefit if selection for resistant cultivars could be realized using genetic markers. For southern anthracnose we found one locus on chromosome 1 explaining more than 16.8% of the variation in resistance to single-spore inoculation and 14.3% of the variation in resistance to the mixed-spore inoculation. While the genetic basis of southern anthracnose resistance in red clover is largely unknown, resistance to *Colletotrichum* spp. has been studied in other legume species including soybean, common bean, and the model species *M. truncatula* (Ameline-Torregrosa et al. 2008; Yang et al. 2008).

In alfalfa resistance to southern anthracnose is characterized by a strong hypersensitive response, typical for effector-triggered immunity or race-specific resistance (Mould et al. 1991). Consequently, three different *C. trifolii* races (1, 2, 4) and two resistance genes (*An1, An2*) have been described (Elgin and O’Neill 1988; O’Neill 1989; Mould et al. 1991). Elgin and Ostazeski (1985) stated that resistance to race 1 in the tetraploid cultivar ‘Arc’ is induced by a single dominant gene *An1* that is tetrasomically inherited. *An2* conferred resistance to races 1 and 2 in the cultivar ‘Saranac AR’. It is generally accepted that the two genes act independently and are not linked but the effect of *An1* can be masked by the presence of *An2* (Elgin and Ostazeski 1985; O’Neill 1989). Unfortunately, the model of single tetrasomic gene inheritance could not be verified using Australian germplasm. Depending on the plant material, resistance was either simply inherited and of qualitative nature or of quantitative nature where several QTL with small-to-medium effects were involved in resistance (Mackie et al. 2003, 2007; Irwin et al. 2006).

In *M. truncatula*, where extensive genomic and genetic resources are available, a major QTL on linkage group (LG) 4 governed resistance to *C. trifolii* race 1 and race 2, while a minor QTL on LG6 was only found when inoculated with *C. trifolii* race 1. The QTL on LG4 explained about 40% of the total phenotypic variation and contained a cluster of *NLR* genes (Ameline-Torregrosa et al. 2008). A single dominant gene named *RCT1* on chromosome 4 was mapped in an F2 population that conferred resistance to race 1 (Yang et al. 2008). The *RCT1* gene of *M. truncatula* was transferred into susceptible alfalfa plants. The alfalfa plants carrying the *RCT1* gene from *M. truncatula* were resistant to all three *C. trifolii* races (Yang et al. 2008). Orthologs of *RCT1* were identified in the red clover reference genome sequence using BLASTn, but these genes were not in the flanking regions of any SNPs significantly associated with southern anthracnose resistance in the red clover EUCLEG-accessions.

Nevertheless, the significantly associated SNPs and their 10 kb flanking regions did contain orthologs of putative resistance genes of *M. truncatula*. One SNP explaining 16.8% of the variation to southern anthracnose resistance is located in the red clover gene *Tp57577_TGAC_v2_gene4880*. Its *M. truncatula* ortholog *MTR_1g10302* plays a role in fatty acid synthesis and its *A. thaliana* ortholog, known as *KASI*, is involved in lipid metabolism and plays a role in cell structure and several plant developmental processes (Wu and Xue 2010). Pathways controlling fatty acid metabolism can play significant roles in cuticular plant defense (Kachroo and Kachroo 2009), hence we speculate that the gene *Tp57577_TGAC_v2_gene4880* might be involved in resistance to southern anthracnose.

We found eight SNPs that were significantly associated with clover rot resistance, explaining together 80.2% of the total phenotypic variation (Table 2). Our study is the first to report loci associated to *S. trifoliorum* resistance in red clover. QTL for resistance to the related species *S. sclerotiorum* have been identified in legume crops such as soybean (Kim and Diers 2000; Arahana et al. 2001), and common bean (Park et al. 2001; Miklas 2007). Most of these QTL explained between 10% and 23% of the variation, which is comparable to the QTL identified in our study. The significant SNPs found for clover rot resistance are located in genes on chromosomes 1, and in scaffolds not assigned to chromosomes. The orthologs of the gene *Tp57577_TGAC_v2_gene37747* and the gene *Tp57577_TGAC_v2_gene968* are known in *M. truncatula* to be putative resistance genes. These putative disease resistance genes encode the RPP13-like protein, which in *Arabidopsis* has been reported to be involved in protection of plants against pathogen invasion by triggering a specific defense system against downy mildew (Bittner-Eddy et al. 2000). Resistance to clover rot in red clover is widely assumed to be a quantitative trait (Poland et al. 2009; Klimenko et al. 2010; Vleugels and van Bockstaele 2013). For quantitative resistance traits, MAS is often ineffective due to population-specific effects and the lack of validation in unrelated populations (Miedaner and Korzun 2012). Therefore, MAS is most likely not effective enough to successfully replace artificial inoculation to select for clover rot resistance. Genomic selection might be a more promising approach (Miedaner et al. 2020). Based on literature (Mackie et al. 2007; Yang et al. 2008) and since only one SNP explained a substantial portion of the variation, we assume that inheritance of southern anthracnose resistance in red clover is mono-or oligogenic. If the effect of the QTL found here can be confirmed, implementing MAS in red clover breeding programs will become a feasible strategy to improve southern anthracnose resistance.

We conducted association studies on allele frequencies per population, allowing to reveal the high genetic population variation of outcrossing species such as red clover, without the need to sequence thousands of individuals (Byrne et al. 2013). If sequencing resources are limited, association studies on population level provide a valuable method and have been effective in finding important loci in humans (Riaz et al. 2016), and to a lesser extent also in plants (Cericola et al. 2018; Keep et al. 2020). Nevertheless, to validate the significant loci and to further characterize resistance alleles, genotyping of single plants is necessary.

Given current and predicted future disease pressure, cultivars that are resistant to southern anthracnose and to clover rot are urgently needed to ensure successful red clover production in Central Europe. Independent inheritance seems likely since Pearson correlation between accession means for the two diseases was absent (0.058, data not shown), and resistance loci found in this study were on different chromosomes (scaffolds) or far apart and are most likely not linked. Therefore, the chances are high that the two diseases are not associated and combining them in one population seems reasonable. We expect that our findings provide a path forward to increase efficiency in breeding for disease resistance in red clover.

## Data availability

Raw data and scripts can be accessed via: https://doi.org/10.5281/zenodo.6572010

## Author contribution

LAF performed the research on southern anthracnose resistance, analyzed all data and drafted the manuscript. TV organized and performed the clover rot trials and contributed to the writing. TR and LS designed the genotyping strategy. LS was responsible for the DNA extraction. TR performed the variant calling and the SNP allele frequency estimation of the pool-GBS data. MP assisted with data analysis. RK, FXS, IRR, BS and CG supported the research design. RK, IRR, and BS helped interpreting the results and drafting the manuscript. All authors read and approved the final version of the manuscript.

## Acknowledgements

We thank Amir Saleem, Philipp Streckeisen and the technical teams at ILVO and Agroscope for their excellent technical support. Many thanks to Sabine Van Glabeke for excellent bio-informatics analyses, Paul Schmidt from the BioMath GmbH for the support on the phenotypic analysis and Daniel Ariza-Suarez of the Molecular Plant Breeding group at ETH Zurich for the valuable discussions on the statistical analysis. We thank all the participants of the EUCLEG project that established the EUCLEG-accession set, and all institutions providing red clover seeds, including: AgResearch (NZ), Agricultural Res. Ltd. (CZ), Agroscope, (CH), Boreal (FI), DLF Seeds (CZ), DSV (DE), Graminor (NO), HBLFA (AT), Hokkaido Ag. Res. (JP), IBERS (UK), IFVCNS (RS), ILVO (BE), INRA (FR), Lantmännen (SE), NordGen (SE), PGG Wrightson (NZ), RAGT2n (FR), and USDA (USA).

## Abbreviation

GBS: Genotyping-by-Sequencing
GWAS: Genome-wide association studies
MAS: Marker-assisted selection

## References

Adam-Blondon AF, Sévignac M, Bannerot H, Dron M (1994) SCAR, RAPD and RFLP markers linked to a dominant gene (Are) conferring resistance to anthracnose in common bean. Theoret Appl Genetics 88:865–870. https://doi.org/10.1007/BF01253998

Ameline-Torregrosa C, Cazaux M, Danesh D, et al (2008) Genetic dissection of resistance to anthracnose and powdery mildew in Medicago truncatula. MPMI 21:61–69. https://doi.org/10.1094/MPMI-21-1-0061

Arahana VS, Graef GL, Specht JE, et al (2001) Identification of QTLs for resistance to Sclerotinia sclerotiorum in soybean. Crop Sci 41:180–188. https://doi.org/10.2135/cropsci2001.411180x

Bain SM, Essary SH (1906) A new anthracnose of alfalfa and red clover. The Journal of Mycology 12:192. https://doi.org/10.2307/3753010

Bittner-Eddy PD, Crute IR, Holub EB, Beynon JL (2000) RPP13 is a simple locus in Arabidopsis thaliana for alleles that specify downy mildew resistance to different avirulence determinants in Peronospora parasitica. Plant J 21:177–188. https://doi.org/10.1046/j.1365-313x.2000.00664.x

Boller B, Posselt UK, Veronesi F (eds) (2010) Fodder crops and amenity grasses. Springer New York, New York, NY

Bonnafous F, Duhnen A, Gody L, et al (2019) mlmm.gwas: Pipeline for GWAS using MLMM. Version R package version 1.0.6URL https://CRAN.R-project.org/package=mlmm.gwas

Broderick GA (1995) Desirable characteristics of forage legumes for improving protein utilization in ruminants. Journal of Animal Science 73:2760. https://doi.org/10.2527/1995.7392760x

Burdon JJ, Thrall PH (2009) Coevolution of plants and their pathogens in natural habitats. Science 324:755–756. https://doi.org/10.1126/science.1171663

Butler DG, Cullis BR, Gilmour AR, et al. (2017) ASReml-R reference manual version 4. VSN International Ltd, Hemel Hempstead, HP1 1ES, UK

Byrne S, Czaban A, Studer B, et al (2013) Genome wide allele frequency fingerprints (GWAFFs) of populations via genotyping by sequencing. PLoS ONE 8:e57438. https://doi.org/10.1371/journal.pone.0057438

Cericola F, Lenk I, Fè D, et al (2018) Optimized use of low-depth genotyping-by-sequencing for genomic prediction among multi-parental family pools and single plants in perennial ryegrass (Lolium perenne L.). Front Plant Sci 9:369. https://doi.org/10.3389/fpls.2018.00369

Chaloner TM, Gurr SJ, Bebber DP (2021) Plant pathogen infection risk tracks global crop yields under climate change. Nat Clim Chang 11:710–715. https://doi.org/10.1038/s41558-021-01104-8

Chen J, Chen Z (2008) Extended Bayesian information criteria for model selection with large model spaces. Biometrika 95:759–771. https://doi.org/10.1093/biomet/asn034

Collard BCY, Mackill DJ (2008) Marker-assisted selection: an approach for precision plant breeding in the twenty-first century. Phil Trans R Soc B 363:557–572. https://doi.org/10.1098/rstb.2007.2170

Cullis BR, Smith AB, Coombes NE (2006) On the design of early generation variety trials with correlated data. JABES 11:381–393. https://doi.org/10.1198/108571106X154443

De Silva DD, Crous PW, Ades PK, et al (2017) Life styles of Colletotrichum species and implications for plant biosecurity. Fungal Biology Reviews 31:155–168. https://doi.org/10.1016/j.fbr.2017.05.001

De Vega JJ, Ayling S, Hegarty M, et al (2015) Red clover (Trifolium pratense L.) draft genome provides a platform for trait improvement. Sci Rep 5:17394. https://doi.org/10.1038/srep17394

Dean R, Van Kan JAL, Pretorius ZA, et al (2012) The Top 10 fungal pathogens in molecular plant pathology: Top 10 fungal pathogens. Molecular Plant Pathology 13:414–430. https://doi.org/10.1111/j.1364-3703.2011.00783.x

Delclos B, Duc G (1996) Etude de la résistance à Sclerotinia trifoliorum chez le trèfle violet (Trifolium pratense L.) Dissertation, University of Paris

Elgin JH, O’Neill NR (1988) Comparison of genes controlling race 1 anthracnose resistance in Arc and Saranac AR alfalfa. Crop Sci 28:657–659. https://doi.org/10.2135/cropsci1988.0011183X002800040020x

Guo Z, Wang H, Tao J, et al (2019) Development of multiple SNP marker panels affordable to breeders through genotyping by target sequencing (GBTS) in maize. Mol Breeding 39:37. https://doi.org/10.1007/s11032-019-0940-4

Halling MA, Topp CFE, Doyle CJ (2004) Aspects of the productivity of forage legumes in Northern Europe. Grass and Forage Sci 59:331–344. https://doi.org/10.1111/j.1365-2494.2004.00435.x

Hartmann S, Schubiger FX, Grieder C, Wosnitza A (2022) A decade of variety testing for resistance of red clover to southern anthracnose (Colletotrichum trifolii Bain et Essary) at the Bavarian state research center for agriculture (LfL). Agriculture 12:. https://doi.org/10.3390/agriculture12020249

Huxley J (1939) Clines: An auxiliary method in taxonomy. Bijdr Dierk 27:491–520

Irwin JAG, Aitken KS, Mackie JM, Musial JM (2006) Genetic improvement of lucerne for anthracnose (Colletotrichum trifolii) resistance. Austral Plant Pathol 35:573. https://doi.org/10.1071/AP06059

Jacob I, Hartmann S, Schubiger FX, Struck C (2015) Resistance screening of red clover cultivars to Colletotrichum trifolii and improving the resistance level through recurrent selection. Euphytica 204:303–310. https://doi.org/10.1007/s10681-014-1323-x

Kachroo A, Kachroo P (2009) Fatty acid–derived signals in plant defense. Annu Rev Phytopathol 47:153–176. https://doi.org/10.1146/annurev-phyto-080508-081820

Keep T, Sampoux J-P, Blanco-Pastor JL, et al (2020) High-throughput genome-wide genotyping to optimize the use of natural genetic resources in the grassland species perennial ryegrass (Lolium perenne L.). G3 Genes|Genomes||Genetics 10:3347–3364. https://doi.org/10.1534/g3.120.401491

Kim HS, Diers BW (2000) Inheritance of partial resistance to Sclerotinia stem rot in soybean. Crop Sci 40:55–61. https://doi.org/10.2135/cropsci2000.40155x

Klimenko I, Razgulayeva N, Gau M, et al (2010) Mapping candidate QTL related to plant persistency in red clover. Theor Appl Genet 120:1253–1263. https://doi.org/10.1007/s00122-009-1253-5

Lundin P, Jönsson HA (1974) Weibull’s Britta - a new medium late diploid red clover with a high resistance to clover rot. Agri Hortique Genetica 32:44–54

Mackie JM, Musial JM, Armour DJ, et al (2007) Identification of QTL for reaction to three races of Colletotrichum trifolii and further analysis of inheritance of resistance in autotetraploid lucerne. Theor Appl Genet 114:1417–1426. https://doi.org/10.1007/s00122-007-0527-z

Mackie JM, Musial JM, O’Neill NR, Irwin JAG (2003) Pathogenic specialisation within Colletotrichum trifolii in Australia, and lucerne cultivar reactions to all known Australian pathotypes. Aust J Agric Res 54:829. https://doi.org/10.1071/AR03079

Marum P, Smith RR, Grau CR (1994) Development of procedures to identify red clover resistant to Sclerotinia trifoliorum. Euphytica 77:257–261. https://doi.org/10.1007/BF02262639

Miedaner T, Boeven ALG-C, Gaikpa DS, et al (2020) Genomics-assisted breeding for quantitative disease resistances in small-grain cereals and maize. IJMS 21:9717. https://doi.org/10.3390/ijms21249717

Miedaner T, Flath K (2007) Effectiveness and environmental stability of quantitative powdery mildew (Blumeria graminis) resistance among winter wheat cultivars. Plant Breeding 126:553–558. https://doi.org/10.1111/j.1439-0523.2006.01353.x

Miedaner T, Korzun V (2012) Marker-assisted selection for disease resistance in wheat and barley breeding. Phytopathology 102:560–566. https://doi.org/10.1094/PHYTO-05-11-0157

Miklas PN (2007) Marker-assisted backcrossing QTL for partial resistance to Sclerotinia white mold in dry bean. Crop Sci 47:935–942. https://doi.org/10.2135/cropsci2006.08.0525

Mould MJR, Boland GJ, Robb J (1991) Ultrastructure of the Colletotrichum trifolii-Medicago sativa pathosystem. II. Post-penetration events. Physiological and Molecular Plant Pathology 38:195–210. https://doi.org/10.1016/S0885-5765(05)80124-9

Nyfeler D, Huguenin-Elie O, Suter M, et al (2011) Grass–legume mixtures can yield more nitrogen than legume pure stands due to mutual stimulation of nitrogen uptake from symbiotic and non-symbiotic sources. Agriculture, Ecosystems & Environment 140:155–163. https://doi.org/10.1016/j.agee.2010.11.022

Öhberg H (2008) Studies of the persistence of red clover cultivars in Sweden: with particular reference to Sclerotinia trifoliorum. Dept. of Agricultural Research for Northern Sweden, Swedish University of Agricultural Sciences

O’Neill NR (1989) Characterization of induced resistance to anthracnose in alfalfa by races, isolates, and species of Colletotrichum. Phytopathology 79:750. https://doi.org/10.1094/Phyto-79-750

Park SO, Coyne DP, Steadman JR, Skroch PW (2001) Mapping of QTL for resistance to white mold disease in common bean. Crop Sci 41:1253–1262. https://doi.org/10.2135/cropsci2001.4141253x

Patella A, Scariolo F, Palumbo F, Barcaccia G (2019) Genetic structure of cultivated varieties of radicchio (Cichorium intybus L.): A comparison between F1 hybrids and synthetics. Plants 8:213. https://doi.org/10.3390/plants8070213

Piepho H-P, Möhring J (2007) Computing heritability and selection response from unbalanced plant breeding trials. Genetics 177:1881–1888. https://doi.org/10.1534/genetics.107.074229

Poland JA, Balint-Kurti PJ, Wisser RJ, et al (2009) Shades of gray: the world of quantitative disease resistance. Trends in Plant Science 14:21–29. https://doi.org/10.1016/j.tplants.2008.10.006

R Core Team (2021). R: A language and environment for statistical computing. R Foundation for Statistical Computing, Vienna, Austria. URL https://www.R-project.org/

Riaz M, Lorés-Motta L, Richardson AJ, et al (2016) GWAS study using DNA pooling strategy identifies association of variant rs4910623 in OR52B4 gene with anti-VEGF treatment response in age-related macular degeneration. Sci Rep 6:37924. https://doi.org/10.1038/srep37924

RStudio Team (2020). RStudio: Integrated Development for R. RStudio, PBC, Boston, MA URL http://www.rstudio.com/

Saharan GS, Mehta N (2010) Sclerotinia diseases of crop plants: biology, ecology and disease management. Springer, Dordrecht

Schubiger F Xaver, Streckeisen P, Boller B (2003) Resistance to southern anthracnose (Colletotrichum trifolii) in cultivars of red clover (Trifolium pratense). Czech Journal of Genetics and Plant Breeding 39:399

Schubiger FX, Alconz E, Streckeisen P, Boller B (2004) Resistenz von Rotklee gegen den südlichen Stängelbrenner. Agrarforschung 11:168–173

Segura V, Vilhjálmsson BJ, Platt A, et al (2012) An efficient multi-locus mixed-model approach for genome-wide association studies in structured populations. Nat Genet 44:825–830. https://doi.org/10.1038/ng.2314

Tang H, Krishnakumar V, Bidwell S, et al (2014) An improved genome release (version Mt4.0) for the model legume Medicago truncatula. BMC Genomics 15:312. https://doi.org/10.1186/1471-2164-15-312

Taylor NL (2008) A Century of clover breeding developments in the United States. Crop Science 48:1–13. https://doi.org/10.2135/cropsci2007.08.0446

Taylor NL, Quesenberry KH (1996) Red clover science. Kluwer Academic Publishers, Dordrecht ; Boston

Terán H, Singh SP (2009) Recurrent selection for physiological resistance to white mould in dry bean: Recurrent selection for physiological resistance to white mould. Plant Breeding 129:327–333. https://doi.org/10.1111/j.1439-0523.2009.01679.x

Therneau TM (2020) Mixed effects Cox models. Version R package version 2.2-16URL https://CRAN.R-project.org/package=coxme

Vleugels T, Baert J, van Bockstaele E (2013a) Morphological and pathogenic characterization of genetically diverse Sclerotinia Isolates from European red clover crops (Trifolium Pratense L.). J Phytopathol 161:254–262. https://doi.org/10.1111/jph.12056

Vleugels T, Cnops G, van Bockstaele E (2013b) Screening for resistance to clover rot (Sclerotinia spp.) among a diverse collection of red clover populations (Trifolium pratense L.). Euphytica 194:371–382. https://doi.org/10.1007/s10681-013-0949-4

Vleugels T, van Bockstaele E (2013) Number of involved genes and heritability of clover rot (Sclerotinia trifoliorum) resistance in red clover (Trifolium pratense). Euphytica 194:137–148. https://doi.org/10.1007/s10681-013-0982-3

Wu G-Z, Xue H-W (2010) Arabidopsis β-ketoacyl-[acyl carrier protein] synthase I is crucial for fatty acid synthesis and plays a role in chloroplast division and embryo development. The Plant Cell 22:3726–3744. https://doi.org/10.1105/tpc.110.075564

Xiao S, Ellwood S, Calis O, et al (2001) Broad-spectrum mildew resistance in Arabidopsis thaliana mediated by RPW8. Science 291:118–120. https://doi.org/10.1126/science.291.5501.118

Yang S, Gao M, Xu C, et al (2008) Alfalfa benefits from Medicago truncatula: The RCT1 gene from M. truncatula confers broad-spectrum resistance to anthracnose in alfalfa. Proceedings of the National Academy of Sciences 105:12164–12169. https://doi.org/10.1073/pnas.0802518105

Zhou F, Kurth J, Wei F, et al (2001) Cell-autonomous expression of barley Mla1 confers race-specific resistance to the powdery mildew fungus via a Rar1 -independent signaling pathway. Plant Cell 13:337–350. https://doi.org/10.1105/tpc.13.2.337

