## Supplementary Figures for "Phenotypic variation and quantitative trait loci for resistance to southern anthracnose and clover rot in red clover"

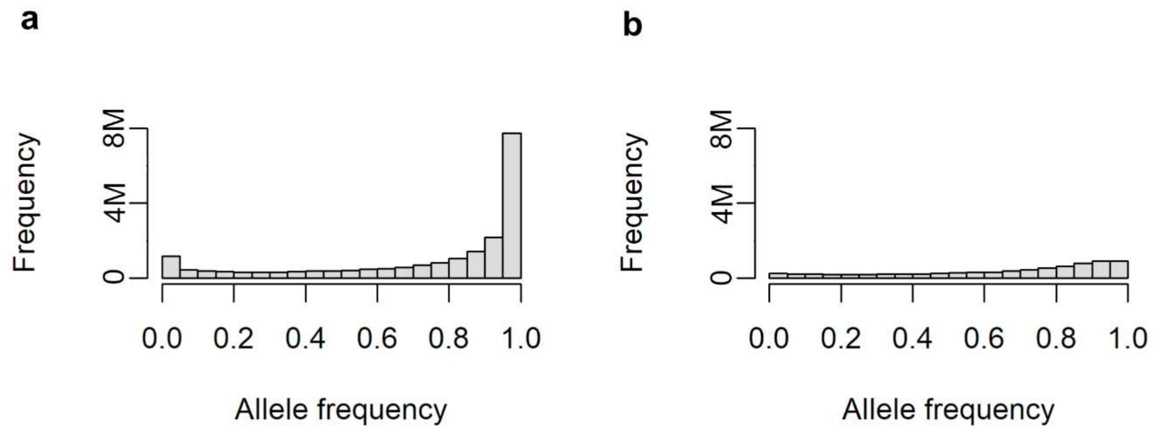


**Supplementary Fig. S1** Allele frequency distribution of all 20,137 single nucleotide polymorphisms (SNPs) before (a) and after (b) filtering. Filtering was performed retaining SNPs with less than 5% missing values, allele frequencies between 0.05 and 0.95 in at least 10 accessions and mean allele frequencies across all accessions between 0.05 and 0.95 (0.05 < MAF < 0.95)

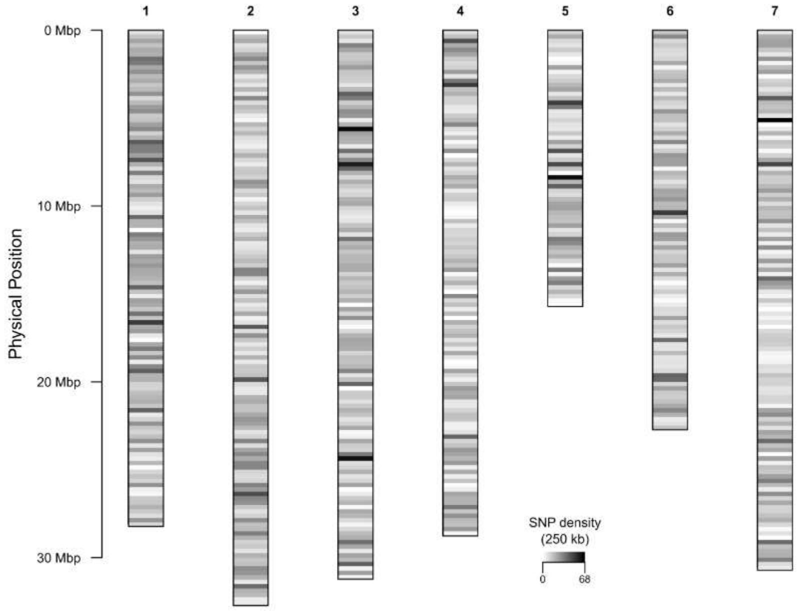


**Supplementary Fig. S2** Single nucleotide polymorphism (SNP) density on the seven *Trifolium pratense* chromosomes. The y-axis represented the interval distance in Mbp. The window size to calculate SNP density was 250kb

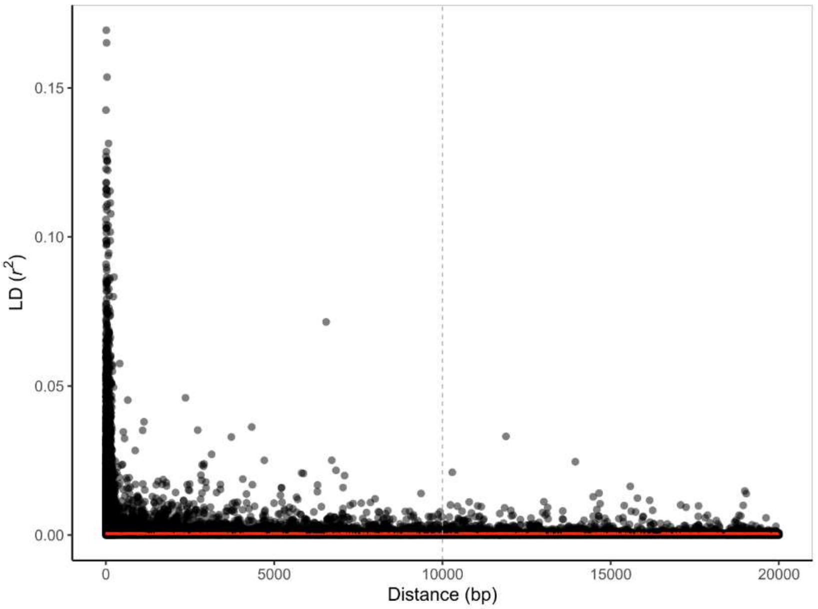


**Supplementary Fig. S3** Linkage disequilibrium (LD) decay against the genetic distance for pairs of SNPs across all seven chromosomes of the red clover genome. LD estimates are reported as squared correlations of allele frequencies ($r^2$). Smoothed fitted lines were calculated using the LOESS method (red solid line). Vertical dotted line is drawn at 10,000 bp
