## Supplementary Methods for "Phenotypic variation and quantitative trait loci for resistance to southern anthracnose and clover rot in red clover"

Genotyping and SNP filtering

Single leaves of 200 plants per accession were pooled and DNA was extracted from the pooled leaf material. All accessions were genotyped by LGC Genomics (Germany) using a *Pst*I-*Mse*I double-digest genotyping-by-sequencing (GBS) method that includes molecular normalization of read depth across loci and size selection of fragments of 100-250 bp (peak around 175 bp) and 2x150 bp Illumina short read sequencing. Reads were demultiplexed with Cutadapt 3.3 (Martin 2011). 3' restriction site remnants, common adapter sequences, and 5' restriction site remnants were removed using a custom python script, Cutadapt 3.3, and FASTX-Toolkit 0.0.14 (Gordon and Hannon 2010). Paired reads were merged with a minimum overlap of 10 bp with PEAR (Zhang et al. 2014). Merged reads were quality filtered and reads shorter than 60 bp were discarded. Reads were aligned to the red clover reference genome sequence v2.1 with the BWA-mem algorithm in BWA 0.7.17 with default parameters (Li 2013; De Vega et al. 2015). Alignments were sorted, indexed and filtered on mapping quality 20 (q20) with SAMtools 1.10 (Li et al. 2009). The Watterson's theta estimator was calculated with NPStat v0.99 (Ferretti et al. 2013). BAM files were converted to mpileup format with SAMtools. Input of NPStat was a mpileup file in which all genome positions with minimum read depth of 30 were concatenated, thus joining the neighboring GBS stacks and excluding the part of the genome without coverage. NPStat was run with the following settings: minor allele counts equal to one read (MAC1), window-size equal to 10,000 bp (i.e. containing about 60 GBS stacks), haploid sample size equal to twice the number of individuals per population, with a maximum of 120 (the maximum number technically accepted in NPStat), and Maximum Coverage equal to 500. Loci with very high (>500) read depth may be derived from repetitive sequences that are mapped onto a single GBS locus and were thus excluded. Per population, a single genome-wide theta value was calculated as the mean across all windows (about 500 windows per sample). The Bayesian SNP calling algorithm implemented in SNAPE-pooled was used to identify SNPs in pool-GBS data (Raineri et al. 2012). SNAPE-pooled was run with settings: - priortype = informative, -fold = folded, -nchr = 120 for consistency with NPStat, as we used the NPStat derived theta values per pool-GBS sample as diversity prior. We used a custom python script to apply filters on the SNAPE-pooled Reference Allele

Frequency (RAF) data. Filters were applied in the following order: (i) SNP positions were deleted if the reference allele was not A, C, G, or T; (ii) SNP frequencies were set to missing data per sample when the two observed alleles were both different from the reference allele, or when the sum of the reference and the alternative allele read counts was lower than 30, (iii) using the Bayesian estimates of the probability of allele presence provided by SNAPE-pooled, we set the alternative allele frequency (and allele counts) to 0 if $p(freq_{alt} \neq 0) < 0.95$ and the reference allele frequency (and allele counts) to 0 if $p(freq_{ref} \neq 0) < 0.95$, (iv) filtered out loci with low coverage (minimal read depth 27) that remained after removing read counts with filter (iii). Next, we integrated all SNP frequency data into one matrix with all samples and all polymorphic loci, and applied filters v-vii: (v) we discarded SNP positions with more than two remaining SNP alleles across all samples (thus removing potential residual low frequency sequencing errors); (vi) we retained only SNPs for which at least 10 accessions had a RAF between 0.05 and 0.95 and the mean allele frequency over all accessions needed to be in the same range; (vii) only SNPs with a maximum of 5% missing values were kept. All missing data points were replaced by the mean allele frequency across all accessions per SNP.

**References supplemental methods:**


De Vega JJ, Ayling S, Hegarty M, et al (2015) Red clover (*Trifolium pratense* L.) draft genome provides a platform for trait improvement. Sci Rep 5:17394. doi: 10.1038/srep17394

Ferretti L, Ramos-Onsins SE, Perez-Enciso M. (2013) Population genomics from pool sequencing. Mol Ecol. 22(22):5561–5576. doi: 10.1111/mec.12522 PMID: 24102736

Gordon A, Hannon G. (2010) Fastx-toolkit. FASTQ/A short-reads preprocessing tools (unpublished).

Li H. (2013) Aligning sequence reads, clone sequences and assembly contigs with BWA-MEM. arXiv:1303.3997v2 [q-bio.GN]

Li H, Handsaker B, Wysoker A, Fennell T, Ruan J, Homer N, et al. (2009). The sequence alignment/map format and SAMtools. Bioinformatics. 25(16):2078–2079. doi :10.1093/bioinformatics/btp352

Martin M. (2011) Cutadapt removes adapter sequences from high-throughput sequencing reads. EMBnet journal. 17(1):pp. 10–12. doi: 10.14806/ej.17.1.200.

Raineri E, Ferretti L, Esteve-Codina A, Nevado B, Heath S, Pérez-Enciso M. (2012) SNP calling by sequencing pooled samples, BMC Bioinformatics,13:239, doi: 10.1186/1471-2105-13-239

Zhang, J., Kobert, K., Flouri, T., & Stamatakis, A. (2014). PEAR: A fast and accurate Illumina Paired End reAd mergeR. Bioinformatics, 30(5), 614–620. doi: 10.1093/bioinformatics/btt593
