## Supplementary Table 1 for "Phenotypic variation and quantitative trait loci for resistance to southern anthracnose and clover rot in red clover"

**Supplementary Table S1** EUCLEG ID, Name of accession, country of origin, year of cultivar registration or collection, type of material as well as collection sites for ecotypes for all accessions used in this study

| EUCLEG ID | Name of accession | Provenance country / Country of origin | Year of cultivar registration / year of collection | Type | Collection site Latitude | Longitude |
|---|---|---|---|---|---|---|
| EUC_TP_001 | Dimanche | FRA | 2006 | Cultivar | | |
| EUC_TP_002 | Discovery | FRA | 2006 | Cultivar | | |
| EUC_TP_003 | Formica | CHE | 1993 | Cultivar | | |
| EUC_TP_004 | Milvus | CHE | 1993 | Cultivar | | |
| EUC_TP_005 | Pavo | CHE | 2002 | Cultivar | | |
| EUC_TP_006 | S586 AberClaret | GBR | 2005 | Cultivar | | |
| EUC_TP_007 | S592 AberChianti | GBR | 2006 | Cultivar | | |
| EUC_TP_008 | Gandalf | NOR | 2015 | Cultivar | | |
| EUC_TP_009 | Lea | NOR | 2002 | Cultivar | | |
| EUC_TP_010 | K 17 | SRB | NA | Cultivar | | |
| EUC_TP_011 | K 38 | SRB | NA | Cultivar | | |
| EUC_TP_012 | SW Ares | SWE | 2001 | Cultivar | | |
| EUC_TP_013 | Cyllene | CZE | 2015 | Cultivar | | |
| EUC_TP_014 | Himalia | CZE | 2012 | Cultivar | | |
| EUC_TP_015 | Metis | CZE | 2016 | Cultivar | | |
| EUC_TP_016 | NA | FRA | 2014 | Cultivar | | |
| EUC_TP_017 | Saija | FIN | 2006 | Cultivar | | |
| EUC_TP_018 | Global | BEL | 2002 | Cultivar | | |
| EUC_TP_019 | Merviot | BEL | 1980 | Cultivar | | |
| EUC_TP_020 | Bonus | CZE | 2008 | Cultivar | | |
| EUC_TP_021 | NGB1132 | FIN | 1981 | Landrace | | |
| EUC_TP_022 | NGB1133 | FIN | 1981 | Landrace | | |
| EUC_TP_023 | NGB1142 | FIN | 1981 | Landrace | | |
| EUC_TP_024 | NGB14322 | FIN | 1998 | Ecotype | NA | NA |
| EUC_TP_025 | NGB1730 | DNK | 1964 | Cultivar | | |
| EUC_TP_026 | NGB2161 | NOR | NA | Landrace | | |
| EUC_TP_027 | NGB2391 | SWE | NA | Landrace | | |
| EUC_TP_028 | NGB2392 | SWE | NA | Landrace | | |
| EUC_TP_029 | NGB2458 | SWE | NA | Landrace | | |

| | | | | |
|---|---|---|---|---|
| EUC_TP_030 | NGB2461 | SWE | NA | Landrace |
| EUC_TP_031 | NGB2487 | SWE | NA | Landrace |
| EUC_TP_032 | NGB2490 | SWE | NA | Landrace |
| EUC_TP_033 | NGB2492 | SWE | NA | Landrace |
| EUC_TP_034 | NGB4089 | SWE | NA | Landrace |
| EUC_TP_035 | Grasslands Colenso | NZL | 1989 | Cultivar |
| EUC_TP_036 | Sensation | NZL | 2002 | Cultivar |
| EUC_TP_037 | Affoltern i.E._328 | CHE | 1972 | Landrace |
| EUC_TP_038 | Affoltern i.E._6 | CHE | 1972 | Landrace |
| EUC_TP_039 | Belpberg_225 | CHE | 1972 | Landrace |
| EUC_TP_040 | Belpberg_226 | CHE | 1972 | Landrace |
| EUC_TP_041 | Belpberg_229 | CHE | 1972 | Landrace |
| EUC_TP_042 | Bern_78 | CHE | 1972 | Landrace |
| EUC_TP_043 | Bowil_119 | CHE | 1972 | Landrace |
| EUC_TP_044 | Bruetten_3 | CHE | 1972 | Landrace |
| EUC_TP_045 | Bubikon_8 | CHE | 1972 | Landrace |
| EUC_TP_046 | Burgistein_300 | CHE | 1972 | Landrace |
| EUC_TP_047 | Columba | CHE | 2016 | Cultivar |
| EUC_TP_048 | Corvus | CHE | 1995 | Cultivar |
| EUC_TP_049 | Dafila | CHE | 2008 | Cultivar |
| EUC_TP_050 | Frauenkappelen_86 | CHE | 1972 | Landrace |
| EUC_TP_051 | Goldbach i.E._167 | CHE | 1972 | Landrace |
| EUC_TP_052 | Huttwil_50 | CHE | 1972 | Landrace |
| EUC_TP_053 | Huttwil_60 | CHE | 1972 | Landrace |
| EUC_TP_054 | Koeniz_231 | CHE | 1972 | Landrace |
| EUC_TP_055 | Koeniz_279 | CHE | 1972 | Landrace |
| EUC_TP_056 | Krauchthal_176 | CHE | 1972 | Landrace |
| EUC_TP_057 | Lanzenhaeusern_291 | CHE | 1972 | Landrace |
| EUC_TP_058 | Lestris | CHE | 2009 | Cultivar |
| EUC_TP_059 | Merula | CHE | 2002 | Cultivar |
| EUC_TP_060 | Milonia | CHE | 2006 | Cultivar |
| EUC_TP_061 | Monaco | CHE | 2012 | Cultivar |
| EUC_TP_062 | Niederwangen_262 | CHE | 1972 | Landrace |
| EUC_TP_063 | Niederwangen_75 | CHE | 1972 | Landrace |
| EUC_TP_064 | Oberthal_121 | CHE | 1972 | Landrace |

| | | | | | | |
|---|---|---|---|---|---|---|
| EUC_TP_065 | Oberuzwil_280 | CHE | 1972 | Landrace | | |
| EUC_TP_066 | Pastor | CHE | 2010 | Cultivar | | |
| EUC_TP_067 | Riedbach_88 | CHE | 1972 | Landrace | | |
| EUC_TP_068 | Rueegsau_160 | CHE | 1972 | Landrace | | |
| EUC_TP_069 | Rueti_314 | CHE | 1972 | Landrace | | |
| EUC_TP_070 | Schmidigen_336 | CHE | 1972 | Landrace | | |
| EUC_TP_071 | Semperina | CHE | 2016 | Cultivar | | |
| EUC_TP_072 | Signau_140 | CHE | 1972 | Landrace | | |
| EUC_TP_073 | Sumiswald_189 | CHE | 1972 | Landrace | | |
| EUC_TP_074 | Ueberstorf_294 | CHE | 1972 | Landrace | | |
| EUC_TP_075 | Ueberstorf_346 | CHE | 1972 | Landrace | | |
| EUC_TP_076 | Ufhusen_52 | CHE | 1972 | Landrace | | |
| EUC_TP_077 | Uttigen_2 | CHE | 1972 | Landrace | | |
| EUC_TP_078 | Wynigen_335 | CHE | 1972 | Landrace | | |
| EUC_TP_079 | Zaeziwil_125 | CHE | 1972 | Landrace | | |
| EUC_TP_080 | Zaeziwil_127 | CHE | 1972 | Landrace | | |
| EUC_TP_081 | Aa 3100 | GBR | 1948 | Cultivar | | |
| EUC_TP_082 | Aa 3148 | GBR | 1948 | Cultivar | | |
| EUC_TP_083 | Aa 3149 | GBR | 1948 | Cultivar | | |
| EUC_TP_084 | AA 32 | GBR | 2011 | Breeding material | | |
| EUC_TP_085 | Aa 3459 | GBR | 1957 | Cultivar | | |
| EUC_TP_086 | Aa 4190 | POL | 1990 | Ecotype | 50.85 | 20.65 |
| EUC_TP_087 | Aa 4292 | CZE | 1992 | Ecotype | 49.483 | 18.1 |
| EUC_TP_088 | Aa 4298 | SVK | 1992 | Ecotype | 49.183 | 18.2167 |
| EUC_TP_089 | Aa 4351 | BGR | 1993 | Ecotype | 42.85 | 25.5667 |
| EUC_TP_090 | Aa 4379 Britta | GBR | 1994 | Cultivar | | |
| EUC_TP_091 | Aa 4390 | PRT | 1995 | Ecotype | 41.333 | -7.7667 |
| EUC_TP_092 | Aa 4400 | GBR | 1996 | Ecotype | 54.65 | -2.15 |
| EUC_TP_093 | Aa 4444 | ITA | 1998 | Ecotype | 46.017 | 12.4833 |
| EUC_TP_094 | Aa 4516 | ESP | 2003 | Ecotype | 43.305 | -4.8352 |
| EUC_TP_095 | Aa 4519 | ESP | 2003 | Ecotype | 42.913 | -5.9247 |
| EUC_TP_097 | Aa 4528 | ESP | 2003 | Ecotype | 43.412 | -5.3078 |
| EUC_TP_098 | Aa 4939 | NZL | 2003 | Cultivar | | |
| EUC_TP_099 | Aa 5674 | ARG | 2016 | Cultivar | | |
| EUC_TP_100 | Aa 5675 | ARG | 2016 | Cultivar | | |

| | | | | |
|---|---|---|---|---|
| EUC_TP_101 | Aa 5676 | ARG | 2016 | Cultivar |
| EUC_TP_102 | Aa 5677 | ARG | 2016 | Cultivar |
| EUC_TP_103 | Aa 5678 | ARG | 2016 | Cultivar |
| EUC_TP_104 | Aa 5746 Harmonie | GBR | 2017 | Cultivar |
| EUC_TP_105 | Aa4380 Altaswede | CAN | NA | Cultivar |
| EUC_TP_106 | S543 AberRuby | GBR | 2000 | Cultivar |
| EUC_TP_108 | TP9525 | CHE | NA | Breeding material |
| EUC_TP_109 | TP9645 | CHE | NA | Breeding material |
| EUC_TP_110 | TP9445 | CHE | NA | Breeding material |
| EUC_TP_111 | TP9735 | CHE | NA | Breeding material |
| EUC_TP_112 | TP9315 | CHE | NA | Breeding material |
| EUC_TP_113 | Gumpensteiner Rotklee | AUT | 1974 | Cultivar |
| EUC_TP_114 | Cinnamon Plus | USA | 2003 | Cultivar |
| EUC_TP_115 | DFRC11 | USA | NA | Breeding material |
| EUC_TP_116 | DFRC12 | USA | NA | Breeding material |
| EUC_TP_117 | DFRC13 | USA | NA | Breeding material |
| EUC_TP_118 | DFRC14 | USA | NA | Breeding material |
| EUC_TP_119 | DFRC15 | USA | NA | Breeding material |
| EUC_TP_120 | FF 9615 | USA | 2013 | Cultivar |
| EUC_TP_121 | Marathon | USA | 1985/1994 | Cultivar |
| EUC_TP_122 | Starfire I | USA | 1999 | Cultivar |
| EUC_TP_123 | Starfire II | USA | 2007 | Cultivar |
| EUC_TP_124 | GnRk0729 | NOR | NA | Breeding material |
| EUC_TP_125 | GnRk0747 | NOR | NA | Breeding material |
| EUC_TP_126 | KvRk0201 | NOR | NA | Breeding material |
| EUC_TP_127 | LGRk8801 | NOR | NA | Breeding material |
| EUC_TP_128 | LGRk9415 | NOR | NA | Breeding material |
| EUC_TP_129 | Linus | NOR | 2014 | Cultivar |
| EUC_TP_130 | LøRk0286 | NOR | NA | Breeding material |
| EUC_TP_131 | LøRk0287 | NOR | NA | Breeding material |
| EUC_TP_132 | Linn | NOR | 2021 | Cultivar |
| EUC_TP_134 | LøRk9207 | NOR | NA | Breeding material |
| EUC_TP_135 | LøRk9625 | NOR | NA | Breeding material |
| EUC_TP_136 | LøRk9627 | NOR | NA | Breeding material |

| | | | | |
|---|---|---|---|---|
| EUC_TP_137 | LøRk9628 | NOR | NA | Breeding material |
| EUC_TP_138 | LøRk9753 | NOR | NA | Breeding material |
| EUC_TP_139 | VåRk0401 | NOR | NA | Breeding material |
| EUC_TP_140 | VåRk0510 | NOR | NA | Breeding material |
| EUC_TP_141 | VåRk0512 | NOR | NA | Breeding material |
| EUC_TP_142 | VåRk0513 | NOR | NA | Breeding material |
| EUC_TP_143 | VåRk0624 | NOR | NA | Breeding material |
| EUC_TP_144 | VåRk0625 | NOR | NA | Breeding material |
| EUC_TP_145 | K 39 | SRB | NA | Cultivar |
| EUC_TP_146 | Diplomat | DEU | 2001 | Cultivar |
| EUC_TP_147 | SW 1479004 | SWE | NA | Breeding material |
| EUC_TP_148 | SW 1578301 | SWE | NA | Breeding material |
| EUC_TP_149 | SW 1678001 | SWE | NA | Breeding material |
| EUC_TP_150 | SW RK1092 | SWE | NA | Breeding material |
| EUC_TP_151 | SW RK1117 | SWE | NA | Breeding material |
| EUC_TP_152 | SW RK1118 | SWE | NA | Breeding material |
| EUC_TP_153 | SW RK1119 | SWE | NA | Breeding material |
| EUC_TP_154 | SW RK1120 | SWE | NA | Breeding material |
| EUC_TP_155 | SW RK1121 | SWE | NA | Breeding material |
| EUC_TP_156 | SW RK1122 | SWE | NA | Breeding material |
| EUC_TP_157 | SW RK1123 | SWE | NA | Breeding material |
| EUC_TP_158 | SW RK1124 | SWE | NA | Breeding material |
| EUC_TP_159 | SW RK1125 | SWE | NA | Breeding material |
| EUC_TP_160 | SW RK1131 | SWE | NA | Breeding material |
| EUC_TP_161 | SW RK1132 | SWE | NA | Breeding material |
| EUC_TP_162 | SW RK1133 | SWE | NA | Breeding material |
| EUC_TP_163 | SW RK1134 | SWE | NA | Breeding material |
| EUC_TP_164 | SW Yngve | SWE | 2005 | Cultivar |
| EUC_TP_165 | SWÅ RK09093 | SWE | NA | Breeding material |
| EUC_TP_166 | 0780MP2 | CZE | 2007 | Breeding material |
| EUC_TP_167 | 08102MP2 | CZE | 2008 | Breeding material |
| EUC_TP_168 | 08102MP4 | CZE | 2008 | Breeding material |
| EUC_TP_169 | Callisto | CZE | 2011 | Cultivar |
| EUC_TP_170 | Elara | CZE | 2014 | Cultivar |
| EUC_TP_171 | Ganymed | CZE | 2016 | Cultivar |

| | | | | |
|---|---|---|---|---|
| EUC_TP_172 | Hegemon | CZE | 2017 | Cultivar |
| EUC_TP_173 | Helike | CZE | 2016 | Cultivar |
| EUC_TP_174 | HŽ 2004 80 - 01 | CZE | 2004 | Breeding material |
| EUC_TP_175 | JL 2n 07 80 MP 1 | CZE | 2007 | Breeding material |
| EUC_TP_176 | Kalyke | CZE | 2017 | Cultivar |
| EUC_TP_177 | TPD-05-11-18002 | CZE | 2011 | Breeding material |
| EUC_TP_178 | TPD-05-11-3087 | CZE | 2011 | Breeding material |
| EUC_TP_179 | TPD-05-11-3088 | CZE | 2011 | Breeding material |
| EUC_TP_180 | TPD-05-13-3080 | CZE | 2013 | Breeding material |
| EUC_TP_181 | TPD-05-13-3085 | CZE | 2013 | Breeding material |
| EUC_TP_182 | TPD-05-13-3091 | CZE | 2013 | Breeding material |
| EUC_TP_183 | TPD-05-14-1011 | CZE | 2014 | Breeding material |
| EUC_TP_184 | TPD-05-15-3127 | CZE | 2015 | Breeding material |
| EUC_TP_185 | TPD-05-15-3128 | CZE | 2015 | Breeding material |
| EUC_TP_186 | TPD-05-15-3129 | CZE | 2015 | Breeding material |
| EUC_TP_187 | TPD-05-16-3076 | CZE | 2016 | Breeding material |
| EUC_TP_188 | TPD-05-16-3146 | CZE | 2016 | Breeding material |
| EUC_TP_189 | TPD-05-16-3177 | CZE | 2016 | Breeding material |
| EUC_TP_190 | NA | FRA | 1986 | Cultivar |
| EUC_TP_191 | NA | FRA | 1984 | Cultivar |
| EUC_TP_192 | NA | FRA | 2012 | Cultivar |
| EUC_TP_193 | NA | FRA | 2010 | Cultivar |
| EUC_TP_194 | Grasslands Hamua | NZL | 1946 | Cultivar |
| EUC_TP_195 | Grasslands Turoa | NZL | 1932 | Cultivar |
| EUC_TP_196 | Relish | NZL | 2012 | Cultivar |
| EUC_TP_197 | Ruby/Enterprise | NZL | ~1990 | Cultivar |
| EUC_TP_198 | Natsuyu | JPN | 2004 | Cultivar |
| EUC_TP_199 | Ryokuyu | JPN | 2012 | Cultivar |
| EUC_TP_200 | NS-Mlava | SRB | 2010 | Cultivar |
| EUC_TP_201 | NS-Petnica | SRB | 2012 | Cultivar |
| EUC_TP_202 | NS-Sana | SRB | 2014 | Cultivar |
| EUC_TP_203 | Una(NS) | SRB | 2016 | Cultivar |
| EUC_TP_204 | Zoja (NS) | SRB | 2012 | Cultivar |
| EUC_TP_205 | Avisto | BEL | 2011 | Cultivar |
| EUC_TP_206 | Crossway | NZL | 2002 | Cultivar |

| | | | | |
|---|---|---|---|---|
| EUC_TP_207 | Lemmon | BEL | 2000 | Cultivar |
| EUC_TP_208 | Merkemse | BEL | 1955 | Landrace |
| EUC_TP_209 | Tp.12.12 | BEL | NA | Breeding material |
| EUC_TP_210 | Tp.14.7 | BEL | NA | Breeding material |
| EUC_TP_211 | Tandy | BEL | 2021 | Cultivar |
| EUC_TP_212 | Agil | CZE | 2009 | Cultivar |
| EUC_TP_214 | Brisk | CZE | 2009 | Cultivar |
| EUC_TP_215 | Chlumecký | CZE | 1935 | Cultivar |
| EUC_TP_218 | Feng | CZE | 2015 | Cultivar |
| EUC_TP_219 | Garant | CZE | 2008 | Cultivar |
| EUC_TP_223 | Respect | CZE | 2010 | Cultivar |
| EUC_TP_224 | Slavín | CZE | 2006 | Cultivar |
| EUC_TP_225 | Slavoj | CZE | 2006 | Cultivar |
| EUC_TP_227 | Spurt | CZE | 2009 | Cultivar |
| EUC_TP_228 | Start | CZE | 1973 | Cultivar |
| EUC_TP_229 | Suez | CZE | 2001 | Cultivar |
| EUC_TP_231 | Trubadur | CZE | 2011 | Cultivar |
| EUC_TP_232 | Van | CZE | 2012 | Cultivar |
| EUC_TP_233 | Vendelín | CZE | 2005 | Cultivar |
| EUC_TP_234 | Vltavín | CZE | 1992 | Cultivar |
| EUC_TP_236 | Zefyr | CZE | 2014 | Cultivar |
| EUC_TP_237 | Harmonie | DEU | 2007 | Cultivar |
| EUC_TP_238 | Regent | DEU | 2008 | Cultivar |
| EUC_TP_239 | NGB1736 | DNK | 1964 | Cultivar |
| EUC_TP_240 | NGB2347 | SWE | 1970 | Cultivar |
| EUC_TP_241 | NGB2349 | SWE | 1941 | Cultivar |
| EUC_TP_242 | NGB2395 | SWE | NA | Landrace |
| EUC_TP_243 | NGB2452 | SWE | NA | Landrace |
| EUC_TP_244 | NGB2453 | SWE | NA | Landrace |
| EUC_TP_245 | NGB2464 | SWE | NA | Landrace |
| EUC_TP_246 | NGB2465 | SWE | NA | Landrace |
| EUC_TP_247 | NGB2466 | SWE | NA | Landrace |
| EUC_TP_248 | NGB2468 | SWE | NA | Landrace |
| EUC_TP_249 | NGB2469 | SWE | NA | Landrace |
| EUC_TP_250 | NGB2471 | SWE | NA | Landrace |

| | | | | |
|---|---|---|---|---|
| EUC_TP_251 | NGB2472 | SWE | NA | Landrace |
| EUC_TP_252 | NGB2473 | SWE | NA | Landrace |
| EUC_TP_253 | NGB2474 | SWE | NA | Landrace |
| EUC_TP_254 | NGB2475 | SWE | NA | Landrace |
| EUC_TP_255 | NGB2476 | SWE | NA | Landrace |
| EUC_TP_256 | NGB2477 | SWE | NA | Landrace |
| EUC_TP_257 | NGB2481 | SWE | NA | Landrace |
| EUC_TP_258 | NGB2482 | SWE | NA | Landrace |
| EUC_TP_259 | NGB2494 | SWE | NA | Landrace |
| EUC_TP_260 | NGB2495 | SWE | NA | Landrace |
| EUC_TP_261 | NGB2569 | SWE | NA | Landrace |
| EUC_TP_262 | NGB2598 | SWE | NA | Landrace |
| EUC_TP_263 | NGB2599 | SWE | NA | Landrace |
| EUC_TP_264 | NGB2600 | SWE | NA | Landrace |
| EUC_TP_265 | NGB2739 | SWE | 1937 | Cultivar |
| EUC_TP_266 | NGB2740 | SWE | 1965 | Cultivar |
| EUC_TP_267 | NGB2742 | SWE | 1957 | Cultivar |
| EUC_TP_268 | NGB2745 | SWE | 1977 | Cultivar |
| EUC_TP_269 | NGB2746 | SWE | 1974 | Cultivar |
| EUC_TP_270 | NGB2747 | SWE | 1970 | Cultivar |
| EUC_TP_271 | NGB2748 | SWE | 1951 | Cultivar |
| EUC_TP_272 | NGB2749 | SWE | 1970 | Cultivar |
| EUC_TP_273 | NGB2750 | SWE | 1960 | Cultivar |
| EUC_TP_274 | NGB2751 | SWE | 1976 | Cultivar |
| EUC_TP_275 | NGB4126 | DNK | 1970 | Cultivar |
| EUC_TP_277 | NGB7510 | SWE | NA | Cultivar |
| EUC_TP_278 | NGB9966 | SWE | NA | Landrace |
| EUC_TP_279 | Diadem | FRA | 2002 | Cultivar |
| EUC_TP_280 | Diper | FRA | 1980 | Cultivar |
| EUC_TP_281 | Diplo | FRA | 2006 | Cultivar |
| EUC_TP_282 | Kindia | FRA | 2006 | Cultivar |
| EUC_TP_283 | Affoltern i.E._325 | CHE | 1972 | Landrace |
| EUC_TP_284 | Bern_76 | CHE | 1972 | Landrace |
| EUC_TP_285 | Bigenthal_163 | CHE | 1972 | Landrace |
| EUC_TP_286 | Biglen_352 | CHE | 1972 | Landrace |

| | | | | | | |
|---|---|---|---|---|---|---|
| EUC_TP_287 | Englisberg_249 | CHE | 1972 | Landrace | | |
| EUC_TP_288 | Grossdietwil_21 | CHE | 1972 | Landrace | | |
| EUC_TP_289 | Haeusernmoos_333 | CHE | 1972 | Landrace | | |
| EUC_TP_290 | Koeniz_239 | CHE | 1972 | Landrace | | |
| EUC_TP_291 | Koeniz_247 | CHE | 1972 | Landrace | | |
| EUC_TP_292 | Lauperswil_138 | CHE | 1972 | Landrace | | |
| EUC_TP_293 | MontCalme | CHE | 1970 | Cultivar | | |
| EUC_TP_294 | Neuenegg_340 | CHE | 1972 | Landrace | | |
| EUC_TP_295 | Niederscherli_273 | CHE | 1972 | Landrace | | |
| EUC_TP_296 | Oberbottigen_7 | CHE | 1972 | Landrace | | |
| EUC_TP_297 | Oberoenz_321 | CHE | 1972 | Landrace | | |
| EUC_TP_298 | Oeschenbach_330 | CHE | 1972 | Landrace | | |
| EUC_TP_299 | Renova | CHE | 1964 | Cultivar | | |
| EUC_TP_300 | Riggisberg_318 | CHE | 1972 | Landrace | | |
| EUC_TP_301 | Rueedisbach_332 | CHE | 1972 | Landrace | | |
| EUC_TP_302 | Rüttinova | CHE | 1984 | Cultivar | | |
| EUC_TP_303 | Schmitten_5 | CHE | 1972 | Landrace | | |
| EUC_TP_304 | Signau_154 | CHE | 1972 | Landrace | | |
| EUC_TP_305 | Wasen i.E._199 | CHE | 1972 | Landrace | | |
| EUC_TP_306 | Weier i.E._327 | CHE | 1972 | Landrace | | |
| EUC_TP_307 | Aa 3090 | GBR | 1948 | Cultivar | | |
| EUC_TP_308 | Aa 3108 | GBR | 1948 | Cultivar | | |
| EUC_TP_310 | Aa 4189 | POL | 1990 | Ecotype | 49.25 | 19.95 |
| EUC_TP_311 | Aa 4297 | CZE | 1992 | Ecotype | 49.483 | 18.2667 |
| EUC_TP_312 | Aa 4398 | GBR | 1996 | Ecotype | 54.683 | -2.2667 |
| EUC_TP_313 | Aa 4403 | GBR | 1996 | Ecotype | 51.233 | -2.6833 |
| EUC_TP_315 | Aa 4445 | ITA | 1998 | Ecotype | 46.05 | 12.8 |
| EUC_TP_316 | Aa 4448 | ITA | 1998 | Ecotype | 45.767 | 13 |
| EUC_TP_317 | Aa 4456 | ITA | 1998 | Ecotype | 45.85 | 12.8 |
| EUC_TP_318 | Aa 4515 | ESP | 2003 | Ecotype | 43.328 | -4.8786 |
| EUC_TP_319 | Aa 4520 | ESP | 2005 | Ecotype | 43.128 | -5.8141 |
| EUC_TP_320 | Aa 4525 | ESP | 2003 | Ecotype | 43.379 | -5.8658 |
| EUC_TP_321 | Aa 4527 | ESP | 2003 | Ecotype | 40.517 | -5.2695 |
| EUC_TP_322 | Aa 4529 | ESP | 2003 | Ecotype | 43.276 | -5.8084 |
| EUC_TP_324 | Aa 4593 | GBR | 2009 | Cultivar | | |

| | | | | |
|---|---|---|---|---|
| EUC_TP_325 | Aa 4934 | GBR | 2012 | Breeding material |
| EUC_TP_326 | Aa 4936 | GBR | 2012 | Breeding material |
| EUC_TP_327 | Aa 4937 | GBR | 2012 | Breeding material |
| EUC_TP_328 | Aa 5417 | GBR | 2015 | Breeding material |
| EUC_TP_329 | SW 1479001 | SWE | NA | Breeding material |
| EUC_TP_330 | SW 1479002 | SWE | NA | Breeding material |
| EUC_TP_331 | SW 1479003 | SWE | NA | Breeding material |
| EUC_TP_332 | SW 1678002 | SWE | NA | Breeding material |
| EUC_TP_333 | SW 1678003 | SWE | NA | Breeding material |
| EUC_TP_334 | SW 1678004 | SWE | NA | Breeding material |
| EUC_TP_335 | SW 1678401 | SWE | NA | Breeding material |
| EUC_TP_336 | SW 1678402 | SWE | NA | Breeding material |
| EUC_TP_337 | SW 1678403 | SWE | NA | Breeding material |
| EUC_TP_338 | SW RK1095 | SWE | NA | Breeding material |
| EUC_TP_339 | SW RK1096 | SWE | NA | Breeding material |
| EUC_TP_340 | SW RK1097 | SWE | NA | Breeding material |
| EUC_TP_341 | SW RK1102 | SWE | NA | Breeding material |
| EUC_TP_342 | SW RK1160 | SWE | NA | Breeding material |
| EUC_TP_343 | SW RK1161 | SWE | NA | Breeding material |
| EUC_TP_344 | SW RK1162 | SWE | NA | Breeding material |
| EUC_TP_345 | SW RK1164 | SWE | NA | Breeding material |
| EUC_TP_346 | SWA 1376104 | SWE | NA | Breeding material |
| EUC_TP_347 | SWA 1376105 | SWE | NA | Breeding material |
| EUC_TP_348 | SWA 1476014 | SWE | NA | Breeding material |
| EUC_TP_349 | SWA 1476016 | SWE | NA | Breeding material |
| EUC_TP_350 | SWA 1476017 | SWE | NA | Breeding material |
| EUC_TP_351 | SWA 1476019 | SWE | NA | Breeding material |
| EUC_TP_352 | SWA 1575301 | SWE | NA | Breeding material |
| EUC_TP_353 | SWA 1575302 | SWE | NA | Breeding material |
| EUC_TP_354 | SWA 1575304 | SWE | NA | Breeding material |
| EUC_TP_355 | SWA 1575305 | SWE | NA | Breeding material |
| EUC_TP_356 | SWA 1575306 | SWE | NA | Breeding material |
| EUC_TP_357 | SWA 1575307 | SWE | NA | Breeding material |
| EUC_TP_358 | SWA 1575308 | SWE | NA | Breeding material |
| EUC_TP_359 | SWA 1575309 | SWE | NA | Breeding material |

| | | | | | | |
|---|---|---|---|---|---|---|
| EUC_TP_360 | SWA 1575312 | SWE | NA | Breeding material | | |
| EUC_TP_361 | SWA 1576005 | SWE | NA | Breeding material | | |
| EUC_TP_362 | SWA 1675205 | SWE | NA | Breeding material | | |
| EUC_TP_363 | SWA 1675206 | SWE | NA | Breeding material | | |
| EUC_TP_364 | SWA 1675207 | SWE | NA | Breeding material | | |
| EUC_TP_365 | SWA 1675208 | SWE | NA | Breeding material | | |
| EUC_TP_366 | SWA 1675212 | SWE | NA | Breeding material | | |
| EUC_TP_367 | TPD-05-04-3000 | CZE | 2004 | Breeding material | | |
| EUC_TP_368 | TPD-05-11-3007 | CZE | 2011 | Breeding material | | |
| EUC_TP_369 | TPD-05-12-3018 | CZE | 2012 | Breeding material | | |
| EUC_TP_370 | Avala (NS) | SRB | 2008 | Cultivar | | |
| EUC_TP_371 | BL-1-Banja Luka | SRB | 1996 | Breeding material | | |
| EUC_TP_372 | BL-3-Banja Luka | SRB | 1996 | Breeding material | | |
| EUC_TP_373 | BL-4-Banja Luka | SRB | 1996 | Breeding material | | |
| EUC_TP_374 | BL-5-Banja Luka | SRB | 1996 | Breeding material | | |
| EUC_TP_375 | D-1 | SRB | 2008 | Breeding material | | |
| EUC_TP_376 | D-10 | SRB | 2008 | Breeding material | | |
| EUC_TP_377 | D-2 | SRB | 2008 | Breeding material | | |
| EUC_TP_378 | D-3 | SRB | 2008 | Breeding material | | |
| EUC_TP_379 | D-4 | SRB | 2008 | Breeding material | | |
| EUC_TP_380 | D-5 | SRB | 2008 | Breeding material | | |
| EUC_TP_381 | D-6 | SRB | 2008 | Breeding material | | |
| EUC_TP_382 | D-7 | SRB | 2008 | Breeding material | | |
| EUC_TP_383 | D-8 | SRB | 2008 | Breeding material | | |
| EUC_TP_384 | D-9 | SRB | 2008 | Breeding material | | |
| EUC_TP_385 | M10-Kopaonik | SRB | 1996 | Ecotype | NA | NA |
| EUC_TP_386 | M11-Kopaonik | SRB | 1996 | Ecotype | NA | NA |
| EUC_TP_387 | M12-Kopaonik | SRB | 1996 | Ecotype | NA | NA |
| EUC_TP_388 | M13-Kopaonik | SRB | 1996 | Ecotype | NA | NA |
| EUC_TP_389 | M14-Kopaonik | SRB | 1996 | Ecotype | NA | NA |
| EUC_TP_390 | NS-Ravanica | SRB | 2010 | Cultivar | | |
| EUC_TP_391 | Broadway | NZL | 2002 | Cultivar | | |
| EUC_TP_392 | Kontiki | DEU | 2010 | Cultivar | | |
| EUC_TP_393 | Mercury | BEL | 1994 | Cultivar | | |
| EUC_TP_394 | Merian | BEL | 2000 | Cultivar | | |

| | | | | |
|---|---|---|---|---|
| EUC_TP_395 | Oudenaerdse | BEL | 1950 | Landrace |
| EUC_TP_396 | Primus | BEL | 1967 | Landrace |
| EUC_TP_397 | Tp.08.4 | BEL | NA | Breeding material |
| EUC_TP_398 | Tp.08.5 | BEL | NA | Breeding material |
| EUC_TP_399 | Violetta | BEL | 1954 | Cultivar |
| EUC_TP_400 | Waesse | BEL | 1950 | Landrace |
| EUC_TP_446 | Affoltern i.E._186 | CHE | 1972 | Landrace |
| EUC_TP_447 | Arni b.Biglen_351 | CHE | 1972 | Landrace |
| EUC_TP_449 | Horgen_1 | CHE | 1972 | Landrace |
| EUC_TP_454 | Lanzenhaeusern_292 | CHE | 1972 | Landrace |
| EUC_TP_456 | Riggisberg_311 | CHE | 1972 | Landrace |
| EUC_TP_495[a] | TP1035 | CHE | 2010 | Breeding material |
| EUC_TP_496[a] | TP1115 | CHE | 2011 | Breeding material |
| EUC_TP_497[a] | TP1135 | CHE | 2011 | Breeding material |
| EUC_TP_498[a] | TP1205 | CHE | 2012 | Breeding material |
| EUC_TP_499[a] | TP1305 | CHE | 2013 | Breeding material |
| EUC_TP_660 | LøRk0498 | NOR | NA | Breeding material |
| EUC_TP_661 | SWA 1575303 | SWE | NA | Breeding material |
| EUC_TP_662 | SWA 1576001 | SWE | NA | Breeding material |

[a] not included in the clover rot resistance trial
