## Supplementary Table 2 for "Phenotypic variation and quantitative trait loci for resistance to southern anthracnose and clover rot in red clover"

**Supplementary Table 2** Accession ID, mean survival rate (%) for southern anthracnose single-spore inoculation, cumulative survival rate (%) for southern anthracnose mixed-spore inoculation and mean resistance index (%) for cover rot as well as standard error (SE) for the three traits and all accessions involved in this study

| EUCLEG ID | Southern anthracnose mean survivor rate (%) single-spore inoculation | SE single-spore inoculation | Southern anthracnose mean cumulative survival rate (%) mixed-spore inoculation[ab] | SE mixed-spore inoculation[a] | Clover rot mean resistance index (%) | SE of resistance index |
|---|---|---|---|---|---|---|
| EUC_TP_001 | 16.84 | 5.61 | 10.92 | 0.67 | 33.72 | 3.37 |
| EUC_TP_002 | 9.87 | 5.62 | 2.63 | 0.67 | 37.31 | 3.34 |
| EUC_TP_003 | 16.29 | 5.61 | 8.48 | 0.67 | 35.71 | 3.35 |
| EUC_TP_004 | 20.61 | 0.96 | 8.72 | 0.11 | 36.72 | 3.24 |
| EUC_TP_005 | 43.66 | 0.96 | 29.93 | 0.11 | 29.07 | 3.35 |
| EUC_TP_006 | 17.40 | 5.62 | 9.18 | 0.67 | 36.26 | 3.35 |
| EUC_TP_007 | 22.25 | 5.62 | 13.80 | 0.67 | 33.91 | 3.35 |
| EUC_TP_008 | 20.64 | 5.62 | 6.14 | 0.67 | 39.48 | 3.38 |
| EUC_TP_009 | 13.82 | 5.62 | 7.08 | 0.67 | 41.46 | 3.35 |
| EUC_TP_010 | 18.98 | 6.48 | 1.31 | 0.78 | 25.02 | 3.35 |
| EUC_TP_011 | 14.55 | 5.62 | 4.21 | 0.67 | 27.07 | 3.40 |
| EUC_TP_012 | 23.55 | 5.62 | 12.60 | 0.67 | 40.32 | 3.35 |
| EUC_TP_013 | 21.82 | 5.62 | 14.52 | 0.67 | 39.77 | 3.34 |
| EUC_TP_014 | 57.19 | 5.62 | 49.77 | 0.67 | 35.19 | 3.36 |
| EUC_TP_015 | 20.03 | 5.62 | 13.53 | 0.67 | 37.75 | 3.35 |
| EUC_TP_016 | 29.90 | 5.62 | 17.00 | 0.67 | 29.32 | 3.34 |
| EUC_TP_017 | 12.95 | 5.62 | 3.88 | 0.67 | 36.14 | 3.35 |
| EUC_TP_018 | 39.17 | 5.62 | 22.59 | 0.67 | 34.35 | 3.16 |
| EUC_TP_019 | 17.80 | 5.62 | 4.70 | 0.67 | 37.45 | 3.36 |
| EUC_TP_020 | 42.89 | 5.61 | 30.18 | 0.67 | 33.24 | 3.34 |
| EUC_TP_021 | 12.20 | 6.48 | 3.81 | 0.78 | 39.84 | 3.39 |
| EUC_TP_022 | 4.95 | 5.62 | 1.77 | 0.67 | 35.66 | 3.25 |
| EUC_TP_023 | 11.32 | 5.62 | 6.25 | 0.67 | 39.17 | 3.34 |
| EUC_TP_024 | 30.69 | 6.48 | 16.64 | 0.78 | 45.91 | 3.27 |
| EUC_TP_025 | 19.54 | 5.62 | 9.15 | 0.67 | 34.21 | 3.34 |
| EUC_TP_026 | 14.09 | 5.62 | 7.02 | 0.67 | 35.66 | 3.34 |
| EUC_TP_027 | 23.18 | 5.62 | 9.03 | 0.67 | 32.99 | 3.37 |
| EUC_TP_028 | 13.24 | 5.61 | 4.18 | 0.67 | 42.33 | 3.41 |
| EUC_TP_029 | 20.50 | 5.62 | 5.81 | 0.67 | 28.68 | 3.43 |
| EUC_TP_030 | 13.05 | 5.62 | 3.96 | 0.67 | 34.37 | 3.35 |
| EUC_TP_031 | 42.88 | 5.62 | 36.66 | 0.67 | 43.36 | 3.35 |
| EUC_TP_032 | 20.22 | 5.62 | 5.01 | 0.67 | 39.84 | 3.34 |
| EUC_TP_033 | 18.30 | 5.62 | 10.92 | 0.67 | 35.89 | 3.35 |
| EUC_TP_034 | 11.88 | 5.62 | 4.74 | 0.67 | 30.90 | 3.36 |
| EUC_TP_035 | 16.06 | 5.61 | 5.18 | 0.67 | 30.18 | 3.24 |
| EUC_TP_036 | 10.63 | 5.61 | 0.52 | 0.67 | 36.41 | 3.34 |
| EUC_TP_037 | 18.45 | 5.61 | 4.04 | 0.67 | 33.47 | 3.23 |

| | | | | | | |
|---|---|---|---|---|---|---|
| EUC_TP_038 | 14.26 | 5.62 | 2.79 | 0.67 | 29.88 | 3.35 |
| EUC_TP_039 | 7.72 | 5.61 | 1.83 | 0.67 | 34.77 | 3.35 |
| EUC_TP_040 | 9.54 | 5.62 | 4.70 | 0.67 | 27.38 | 3.34 |
| EUC_TP_041 | 8.96 | 5.62 | 1.35 | 0.67 | 32.89 | 3.34 |
| EUC_TP_042 | 11.95 | 5.62 | 1.49 | 0.67 | 31.02 | 3.34 |
| EUC_TP_043 | 7.27 | 5.62 | 1.16 | 0.67 | 28.87 | 3.37 |
| EUC_TP_044 | 19.73 | 5.61 | 10.63 | 0.67 | 29.86 | 3.35 |
| EUC_TP_045 | 34.08 | 5.62 | 20.63 | 0.67 | 30.36 | 3.35 |
| EUC_TP_046 | 17.61 | 5.61 | 7.37 | 0.67 | 30.03 | 3.24 |
| EUC_TP_047 | 52.63 | 5.61 | 34.23 | 0.67 | 34.11 | 3.38 |
| EUC_TP_048 | 20.64 | 5.62 | 10.12 | 0.67 | 32.52 | 3.42 |
| EUC_TP_049 | 49.89 | 5.61 | 44.29 | 0.67 | 38.58 | 3.34 |
| EUC_TP_050 | 4.65 | 5.62 | 0.43 | 0.67 | 26.76 | 3.36 |
| EUC_TP_051 | 14.47 | 5.62 | 4.30 | 0.67 | 30.59 | 3.34 |
| EUC_TP_052 | 8.88 | 5.62 | 0.40 | 0.67 | 27.07 | 3.40 |
| EUC_TP_053 | 5.42 | 5.62 | 0.60 | 0.67 | 28.53 | 3.39 |
| EUC_TP_054 | 11.35 | 5.61 | 1.46 | 0.67 | 30.99 | 3.35 |
| EUC_TP_055 | 9.06 | 5.61 | 1.37 | 0.67 | 29.38 | 3.36 |
| EUC_TP_056 | 4.47 | 5.62 | 2.05 | 0.67 | 28.06 | 3.34 |
| EUC_TP_057 | 18.59 | 5.62 | 9.62 | 0.67 | 27.31 | 3.35 |
| EUC_TP_058 | 52.02 | 5.61 | 41.64 | 0.67 | 30.89 | 3.26 |
| EUC_TP_059 | 36.32 | 5.61 | 27.36 | 0.67 | 35.00 | 3.36 |
| EUC_TP_060 | 53.84 | 5.62 | 35.75 | 0.67 | 31.12 | 3.35 |
| EUC_TP_061 | 9.96 | 5.62 | 4.86 | 0.67 | 37.15 | 3.34 |
| EUC_TP_062 | 10.39 | 5.62 | 4.63 | 0.67 | 28.62 | 3.34 |
| EUC_TP_063 | 8.43 | 5.61 | 5.15 | 0.67 | 36.94 | 3.37 |
| EUC_TP_064 | 13.04 | 5.62 | 3.11 | 0.67 | 28.89 | 3.34 |
| EUC_TP_065 | 12.34 | 5.62 | 4.89 | 0.67 | 29.87 | 3.35 |
| EUC_TP_066 | 25.34 | 5.62 | 12.38 | 0.67 | 34.35 | 3.23 |
| EUC_TP_067 | 18.63 | 5.62 | 9.00 | 0.67 | 30.86 | 3.33 |
| EUC_TP_068 | 11.32 | 5.62 | 3.32 | 0.67 | 25.58 | 3.36 |
| EUC_TP_069 | 17.60 | 5.62 | 0.72 | 0.67 | 32.59 | 3.35 |
| EUC_TP_070 | 14.08 | 5.62 | 10.45 | 0.67 | 29.34 | 3.34 |
| EUC_TP_071 | 57.19 | 5.61 | 39.51 | 0.67 | 41.87 | 3.35 |
| EUC_TP_072 | 9.69 | 5.61 | 3.29 | 0.67 | 33.55 | 3.35 |
| EUC_TP_073 | 6.30 | 5.61 | 1.70 | 0.67 | 26.60 | 3.34 |
| EUC_TP_074 | 9.71 | 5.62 | 1.71 | 0.67 | 37.14 | 3.37 |
| EUC_TP_075 | 15.19 | 5.61 | 2.59 | 0.67 | 33.24 | 3.34 |
| EUC_TP_076 | 8.09 | 5.62 | 4.04 | 0.67 | 24.86 | 3.34 |
| EUC_TP_077 | 5.60 | 5.62 | 4.35 | 0.67 | 33.06 | 3.36 |
| EUC_TP_078 | 13.02 | 5.62 | 2.99 | 0.67 | 26.53 | 3.34 |
| EUC_TP_079 | 8.59 | 5.62 | 1.72 | 0.67 | 24.96 | 3.37 |
| EUC_TP_080 | 8.49 | 5.62 | 1.93 | 0.67 | 26.84 | 3.35 |
| EUC_TP_081 | 10.94 | 5.62 | 0.30 | 0.67 | 30.70 | 3.39 |
| EUC_TP_082 | 5.22 | 5.62 | 1.19 | 0.67 | 22.18 | 3.34 |
| EUC_TP_083 | 5.21 | 5.62 | 0.00 | 0.67 | 28.43 | 3.34 |
| EUC_TP_084 | 12.51 | 5.62 | 0.54 | 0.67 | 41.95 | 3.25 |

| | | | | | | |
|---|---|---|---|---|---|---|
| EUC_TP_085 | 13.97 | 5.62 | 5.57 | 0.67 | 25.25 | 3.36 |
| EUC_TP_086 | 10.30 | 5.62 | 0.18 | 0.67 | 30.08 | 3.34 |
| EUC_TP_087 | 46.33 | 5.62 | 35.93 | 0.67 | 30.34 | 3.35 |
| EUC_TP_088 | 17.87 | 5.61 | 4.42 | 0.67 | 33.86 | 3.36 |
| EUC_TP_089 | 3.89 | 5.62 | 0.00 | 0.67 | 30.97 | 3.35 |
| EUC_TP_090 | 26.62 | 5.62 | 20.65 | 0.67 | 38.56 | 3.35 |
| EUC_TP_091 | 2.56 | 5.61 | 0.66 | 0.67 | 23.46 | 3.35 |
| EUC_TP_092 | 5.86 | 5.61 | 0.01 | 0.67 | 33.77 | 3.34 |
| EUC_TP_093 | 21.46 | 5.62 | 18.55 | 0.67 | 41.89 | 3.23 |
| EUC_TP_094 | 19.92 | 5.62 | 5.70 | 0.67 | 25.36 | 3.40 |
| EUC_TP_095 | 16.36 | 5.61 | 11.01 | 0.67 | 26.76 | 3.35 |
| EUC_TP_097 | 6.78 | 5.62 | 4.88 | 0.67 | 26.74 | 3.35 |
| EUC_TP_098 | 26.51 | 5.62 | 14.30 | 0.67 | 27.88 | 3.35 |
| EUC_TP_099 | 58.06 | 5.62 | 42.47 | 0.67 | 31.93 | 3.35 |
| EUC_TP_100 | 57.19 | 5.62 | 49.70 | 0.67 | 32.66 | 3.34 |
| EUC_TP_101 | 12.74 | 5.62 | 4.85 | 0.67 | 23.03 | 3.26 |
| EUC_TP_102 | 42.38 | 5.62 | 34.03 | 0.67 | 32.63 | 3.24 |
| EUC_TP_103 | 40.62 | 5.62 | 32.50 | 0.67 | 31.97 | 3.35 |
| EUC_TP_104 | 18.21 | 5.62 | 8.54 | 0.67 | 41.15 | 3.34 |
| EUC_TP_105 | 12.82 | 5.61 | 4.78 | 0.67 | 32.67 | 3.34 |
| EUC_TP_106 | 12.63 | 5.62 | 0.62 | 0.67 | 28.71 | 3.25 |
| EUC_TP_108 | 17.00 | 5.62 | 3.75 | 0.67 | 37.62 | 3.23 |
| EUC_TP_109 | 31.27 | 5.62 | 18.88 | 0.67 | 37.97 | 3.34 |
| EUC_TP_110 | 45.66 | 5.61 | 30.98 | 0.67 | 36.54 | 3.36 |
| EUC_TP_111 | 52.97 | 5.62 | 43.74 | 0.67 | 37.21 | 3.24 |
| EUC_TP_112 | 38.11 | 5.61 | 23.60 | 0.67 | 42.06 | 3.27 |
| EUC_TP_113 | 8.88 | 5.62 | 0.33 | 0.67 | 28.18 | 3.40 |
| EUC_TP_114 | 73.93 | 5.62 | 65.09 | 0.67 | 43.69 | 3.35 |
| EUC_TP_115 | 51.57 | 5.62 | 44.25 | 0.67 | 21.22 | 3.34 |
| EUC_TP_116 | 70.41 | 5.62 | 57.51 | 0.67 | 30.23 | 3.23 |
| EUC_TP_117 | 56.11 | 5.62 | 48.87 | 0.67 | 31.13 | 3.34 |
| EUC_TP_118 | 57.42 | 5.62 | 49.35 | 0.67 | 32.47 | 3.37 |
| EUC_TP_119 | 67.25 | 5.62 | 45.04 | 0.67 | 32.30 | 3.34 |
| EUC_TP_120 | 53.38 | 5.62 | 42.28 | 0.67 | 35.83 | 3.23 |
| EUC_TP_121 | 50.79 | 6.48 | 43.04 | 0.78 | 26.35 | 3.34 |
| EUC_TP_122 | 56.01 | 5.62 | 49.82 | 0.67 | 30.29 | 3.34 |
| EUC_TP_123 | 62.57 | 5.62 | 55.96 | 0.67 | 29.19 | 3.34 |
| EUC_TP_124 | 29.73 | 5.62 | 18.62 | 0.67 | 40.25 | 3.33 |
| EUC_TP_125 | 11.25 | 5.62 | 6.21 | 0.67 | 44.17 | 3.34 |
| EUC_TP_126 | 20.78 | 5.62 | 12.12 | 0.67 | 37.08 | 3.36 |
| EUC_TP_127 | 33.21 | 5.62 | 28.56 | 0.67 | 37.23 | 3.37 |
| EUC_TP_128 | 27.01 | 5.61 | 15.00 | 0.67 | 41.49 | 3.35 |
| EUC_TP_129 | 16.94 | 5.62 | 9.83 | 0.67 | 29.62 | 3.23 |
| EUC_TP_130 | 39.52 | 5.62 | 27.47 | 0.67 | 37.54 | 3.35 |
| EUC_TP_131 | 16.27 | 5.62 | 4.62 | 0.67 | 33.31 | 3.35 |
| EUC_TP_132 | 29.73 | 5.62 | 18.79 | 0.67 | 34.30 | 3.35 |
| EUC_TP_134 | 19.84 | 5.62 | 12.30 | 0.67 | 39.70 | 3.34 |

| | | | | | | |
|---|---|---|---|---|---|---|
| EUC_TP_135 | 37.24 | 5.62 | 30.34 | 0.67 | 41.67 | 3.37 |
| EUC_TP_136 | 16.55 | 5.61 | 6.56 | 0.67 | 30.51 | 3.34 |
| EUC_TP_137 | 20.66 | 5.62 | 8.76 | 0.67 | 43.04 | 3.37 |
| EUC_TP_138 | 29.70 | 5.62 | 17.51 | 0.67 | 38.51 | 3.34 |
| EUC_TP_139 | 45.83 | 5.61 | 25.32 | 0.67 | 40.11 | 3.34 |
| EUC_TP_140 | 24.29 | 5.62 | 9.98 | 0.67 | 28.17 | 3.35 |
| EUC_TP_141 | 32.90 | 5.62 | 21.33 | 0.67 | 31.74 | 3.23 |
| EUC_TP_142 | 31.77 | 5.61 | 19.14 | 0.67 | 38.08 | 3.35 |
| EUC_TP_143 | 25.32 | 5.62 | 18.74 | 0.67 | 44.63 | 3.34 |
| EUC_TP_144 | 40.63 | 5.62 | 27.41 | 0.67 | 39.07 | 3.35 |
| EUC_TP_145 | 17.32 | 5.62 | 4.73 | 0.67 | 27.39 | 3.37 |
| EUC_TP_146 | 18.68 | 5.62 | 0.96 | 0.67 | 30.77 | 3.34 |
| EUC_TP_147 | 22.16 | 5.62 | 5.70 | 0.67 | 31.06 | 3.35 |
| EUC_TP_148 | 11.13 | 5.61 | 5.24 | 0.67 | 32.41 | 3.23 |
| EUC_TP_149 | 17.51 | 5.62 | 10.40 | 0.67 | 34.93 | 3.36 |
| EUC_TP_150 | 20.24 | 5.62 | 6.87 | 0.67 | 38.66 | 3.34 |
| EUC_TP_151 | 23.25 | 5.62 | 16.08 | 0.67 | 36.56 | 3.34 |
| EUC_TP_152 | 25.31 | 5.62 | 13.34 | 0.67 | 38.01 | 3.34 |
| EUC_TP_153 | 15.17 | 5.62 | 6.55 | 0.67 | 30.58 | 3.35 |
| EUC_TP_154 | 23.49 | 5.62 | 10.18 | 0.67 | 34.81 | 3.35 |
| EUC_TP_155 | 13.20 | 5.62 | 6.75 | 0.67 | 37.44 | 3.39 |
| EUC_TP_156 | 26.36 | 5.62 | 13.50 | 0.67 | 38.79 | 3.24 |
| EUC_TP_157 | 21.01 | 5.61 | 11.19 | 0.67 | 35.62 | 3.35 |
| EUC_TP_158 | 18.14 | 5.61 | 7.64 | 0.67 | 32.56 | 3.35 |
| EUC_TP_159 | 22.17 | 5.61 | 15.13 | 0.67 | 39.17 | 3.24 |
| EUC_TP_160 | 24.15 | 5.62 | 14.93 | 0.67 | 39.22 | 3.35 |
| EUC_TP_161 | 14.10 | 5.62 | 2.58 | 0.67 | 38.03 | 3.35 |
| EUC_TP_162 | 28.74 | 5.62 | 20.74 | 0.67 | 35.86 | 3.23 |
| EUC_TP_163 | 24.86 | 5.62 | 17.06 | 0.67 | 38.31 | 3.35 |
| EUC_TP_164 | 10.42 | 5.61 | 6.91 | 0.67 | 36.30 | 3.36 |
| EUC_TP_165 | 15.19 | 5.61 | 5.56 | 0.67 | 43.17 | 3.34 |
| EUC_TP_166 | 35.05 | 5.62 | 18.94 | 0.67 | 30.78 | 3.36 |
| EUC_TP_167 | 24.19 | 5.62 | 18.38 | 0.67 | 37.92 | 3.36 |
| EUC_TP_168 | 27.25 | 5.62 | 12.62 | 0.67 | 41.85 | 3.34 |
| EUC_TP_169 | 30.15 | 5.62 | 14.93 | 0.67 | 38.77 | 3.35 |
| EUC_TP_170 | 23.73 | 5.62 | 15.95 | 0.67 | 31.29 | 3.35 |
| EUC_TP_171 | 25.83 | 5.62 | 19.59 | 0.67 | 34.05 | 3.34 |
| EUC_TP_172 | 42.77 | 5.62 | 27.62 | 0.67 | 29.40 | 3.35 |
| EUC_TP_173 | 19.84 | 5.61 | 8.76 | 0.67 | 30.62 | 3.35 |
| EUC_TP_174 | 14.23 | 5.61 | 4.26 | 0.67 | 42.51 | 3.38 |
| EUC_TP_175 | 12.71 | 5.62 | 1.43 | 0.67 | 42.98 | 3.35 |
| EUC_TP_176 | 22.36 | 5.62 | 17.55 | 0.67 | 40.35 | 3.24 |
| EUC_TP_177 | 30.46 | 6.48 | 9.81 | 0.78 | 32.40 | 3.39 |
| EUC_TP_178 | 19.68 | 5.61 | 3.39 | 0.67 | 41.07 | 3.34 |
| EUC_TP_179 | 21.67 | 5.61 | 11.48 | 0.67 | 40.06 | 3.24 |
| EUC_TP_180 | 38.38 | 5.62 | 29.13 | 0.67 | 34.09 | 3.34 |
| EUC_TP_181 | 27.06 | 5.62 | 13.12 | 0.67 | 40.81 | 3.23 |

| | | | | | | |
|---|---|---|---|---|---|---|
| EUC_TP_182 | 55.60 | 5.62 | 31.48 | 0.67 | 42.49 | 3.36 |
| EUC_TP_183 | 17.95 | 5.62 | 6.33 | 0.67 | 30.11 | 3.34 |
| EUC_TP_184 | 50.25 | 5.62 | 45.88 | 0.67 | 44.00 | 3.33 |
| EUC_TP_185 | 42.45 | 5.62 | 22.91 | 0.67 | 32.99 | 3.34 |
| EUC_TP_186 | 38.23 | 5.62 | 25.33 | 0.67 | 43.06 | 3.36 |
| EUC_TP_187 | 46.03 | 5.62 | 31.99 | 0.67 | 34.61 | 3.34 |
| EUC_TP_188 | 38.81 | 5.61 | 31.68 | 0.67 | 38.32 | 3.36 |
| EUC_TP_189 | 31.81 | 5.62 | 17.57 | 0.67 | 29.19 | 3.35 |
| EUC_TP_190 | 20.05 | 5.62 | 9.50 | 0.67 | 30.47 | 3.34 |
| EUC_TP_191 | 21.47 | 5.62 | 9.67 | 0.67 | 39.32 | 3.34 |
| EUC_TP_192 | 17.73 | 5.61 | 6.79 | 0.67 | 37.61 | 3.24 |
| EUC_TP_193 | 16.42 | 5.62 | 9.42 | 0.67 | 36.87 | 3.35 |
| EUC_TP_194 | 11.80 | 5.62 | 4.43 | 0.67 | 27.29 | 3.38 |
| EUC_TP_195 | 9.92 | 5.61 | 0.04 | 0.67 | 35.98 | 3.53 |
| EUC_TP_196 | 22.54 | 5.62 | 16.55 | 0.67 | 34.86 | 3.35 |
| EUC_TP_197 | 27.16 | 5.62 | 11.86 | 0.67 | 29.61 | 3.45 |
| EUC_TP_198 | 41.40 | 5.62 | 32.10 | 0.67 | 38.26 | 3.36 |
| EUC_TP_199 | 29.75 | 5.62 | 20.55 | 0.67 | 37.15 | 3.23 |
| EUC_TP_200 | 38.54 | 5.62 | 27.78 | 0.67 | 29.81 | 3.35 |
| EUC_TP_201 | 40.74 | 5.62 | 27.51 | 0.67 | 30.27 | 3.24 |
| EUC_TP_202 | 40.42 | 5.62 | 32.49 | 0.67 | 31.67 | 3.23 |
| EUC_TP_203 | 39.48 | 5.62 | 19.08 | 0.67 | 27.81 | 3.23 |
| EUC_TP_204 | 38.00 | 5.62 | 19.85 | 0.67 | 31.60 | 3.35 |
| EUC_TP_205 | 38.23 | 5.62 | 28.83 | 0.67 | 28.02 | 3.37 |
| EUC_TP_206 | 23.45 | 5.61 | 18.33 | 0.67 | 23.09 | 3.36 |
| EUC_TP_207 | 27.08 | 5.61 | 17.37 | 0.67 | 36.76 | 3.35 |
| EUC_TP_208 | 10.78 | 5.62 | 1.09 | 0.67 | 31.02 | 3.23 |
| EUC_TP_209 | 37.07 | 5.62 | 23.40 | 0.67 | 24.06 | 3.37 |
| EUC_TP_210 | 14.91 | 5.62 | 7.85 | 0.67 | 34.21 | 3.34 |
| EUC_TP_211 | 30.99 | 5.62 | 19.23 | 0.67 | 42.70 | 3.35 |
| EUC_TP_212 | 72.70 | 5.62 | 63.72 | 0.67 | 36.64 | 3.34 |
| EUC_TP_214 | 16.36 | 5.62 | 1.75 | 0.67 | 33.40 | 3.34 |
| EUC_TP_215 | 14.19 | 6.48 | 2.34 | 0.78 | 33.97 | 3.35 |
| EUC_TP_218 | 51.25 | 5.61 | 34.45 | 0.67 | 35.13 | 3.36 |
| EUC_TP_219 | 39.90 | 5.62 | 22.44 | 0.67 | 35.49 | 3.34 |
| EUC_TP_223 | 38.98 | 5.62 | 27.43 | 0.67 | 32.10 | 3.34 |
| EUC_TP_224 | 17.46 | 6.48 | 5.22 | 0.78 | 36.54 | 3.35 |
| EUC_TP_225 | 15.62 | 5.62 | 5.82 | 0.67 | 27.80 | 3.35 |
| EUC_TP_227 | 39.00 | 5.62 | 25.08 | 0.67 | 29.98 | 3.34 |
| EUC_TP_228 | 28.11 | 5.61 | 15.98 | 0.67 | 30.47 | 3.37 |
| EUC_TP_229 | 12.55 | 5.62 | 6.49 | 0.67 | 34.43 | 3.35 |
| EUC_TP_231 | 37.58 | 6.48 | 28.62 | 0.78 | 38.73 | 3.35 |
| EUC_TP_232 | 55.11 | 5.62 | 53.67 | 0.67 | 36.18 | 3.48 |
| EUC_TP_233 | 8.86 | 5.62 | 3.50 | 0.67 | 35.00 | 3.23 |
| EUC_TP_234 | 10.79 | 5.61 | 2.28 | 0.67 | 30.61 | 3.34 |
| EUC_TP_236 | 27.62 | 5.61 | 16.90 | 0.67 | 38.20 | 3.34 |
| EUC_TP_237 | 30.63 | 5.62 | 11.66 | 0.67 | 37.02 | 3.34 |

| | | | | | | |
|---|---|---|---|---|---|---|
| EUC_TP_238 | 19.06 | 5.62 | 7.92 | 0.67 | 36.46 | 3.35 |
| EUC_TP_239 | 20.53 | 5.62 | 13.22 | 0.67 | 29.03 | 3.37 |
| EUC_TP_240 | 34.69 | 5.62 | 21.29 | 0.67 | 38.52 | 3.35 |
| EUC_TP_241 | 26.80 | 5.62 | 17.96 | 0.67 | 29.46 | 3.35 |
| EUC_TP_242 | 19.27 | 5.62 | 12.56 | 0.67 | 30.95 | 3.35 |
| EUC_TP_243 | 15.60 | 5.62 | 2.71 | 0.67 | 31.64 | 3.35 |
| EUC_TP_244 | 9.84 | 5.62 | 2.58 | 0.67 | 30.52 | 3.34 |
| EUC_TP_245 | 11.87 | 5.62 | 5.76 | 0.67 | 35.69 | 3.34 |
| EUC_TP_246 | 9.45 | 5.61 | 6.01 | 0.67 | 26.99 | 3.36 |
| EUC_TP_247 | 20.45 | 5.61 | 3.60 | 0.67 | 28.12 | 3.23 |
| EUC_TP_248 | 14.46 | 5.62 | 1.38 | 0.67 | 40.70 | 3.34 |
| EUC_TP_249 | 7.76 | 5.62 | 2.59 | 0.67 | 32.96 | 3.35 |
| EUC_TP_250 | 15.74 | 5.61 | 7.88 | 0.67 | 42.26 | 3.34 |
| EUC_TP_251 | 11.44 | 5.62 | 5.23 | 0.67 | 39.40 | 3.34 |
| EUC_TP_252 | 17.01 | 5.61 | 3.93 | 0.67 | 35.65 | 3.34 |
| EUC_TP_253 | 12.77 | 5.62 | 1.80 | 0.67 | 32.91 | 3.24 |
| EUC_TP_254 | 16.62 | 5.62 | 5.65 | 0.67 | 39.60 | 3.34 |
| EUC_TP_255 | 13.65 | 5.62 | 6.13 | 0.67 | 38.42 | 3.35 |
| EUC_TP_256 | 14.55 | 5.62 | 5.97 | 0.67 | 34.63 | 3.23 |
| EUC_TP_257 | 13.41 | 5.62 | 1.99 | 0.67 | 33.54 | 3.34 |
| EUC_TP_258 | 10.03 | 5.62 | 2.59 | 0.67 | 33.23 | 3.34 |
| EUC_TP_259 | 7.94 | 5.62 | 0.45 | 0.67 | 38.18 | 3.35 |
| EUC_TP_260 | 15.81 | 5.62 | 2.28 | 0.67 | 32.38 | 3.23 |
| EUC_TP_261 | 15.81 | 5.62 | 5.25 | 0.67 | 34.45 | 3.35 |
| EUC_TP_262 | 15.95 | 5.62 | 5.54 | 0.67 | 29.37 | 3.23 |
| EUC_TP_263 | 12.84 | 5.62 | 8.10 | 0.67 | 29.22 | 3.23 |
| EUC_TP_264 | 14.55 | 5.62 | 5.75 | 0.67 | 31.01 | 3.33 |
| EUC_TP_265 | 24.66 | 5.62 | 16.40 | 0.67 | 38.77 | 3.34 |
| EUC_TP_266 | 10.71 | 5.62 | 3.11 | 0.67 | 36.86 | 3.23 |
| EUC_TP_267 | 15.74 | 6.48 | 6.84 | 0.78 | 39.86 | 3.24 |
| EUC_TP_268 | 25.85 | 5.62 | 11.30 | 0.67 | 38.12 | 3.34 |
| EUC_TP_269 | 20.64 | 5.61 | 7.88 | 0.67 | 37.98 | 3.25 |
| EUC_TP_270 | 13.08 | 5.62 | 3.70 | 0.67 | 36.76 | 3.28 |
| EUC_TP_271 | 10.54 | 5.62 | 5.78 | 0.67 | 32.42 | 3.33 |
| EUC_TP_272 | 15.45 | 5.62 | 6.36 | 0.67 | 32.64 | 3.34 |
| EUC_TP_273 | 5.57 | 5.62 | 3.79 | 0.67 | 28.96 | 3.35 |
| EUC_TP_274 | 10.45 | 5.62 | 1.04 | 0.67 | 38.44 | 3.25 |
| EUC_TP_275 | 23.35 | 5.62 | 13.08 | 0.67 | 34.42 | 3.38 |
| EUC_TP_277 | 30.36 | 5.61 | 21.00 | 0.67 | 33.16 | 3.36 |
| EUC_TP_278 | 17.32 | 6.48 | 6.22 | 0.78 | 37.89 | 3.35 |
| EUC_TP_279 | 20.45 | 6.48 | 10.60 | 0.78 | 35.96 | 3.36 |
| EUC_TP_280 | 20.38 | 5.61 | 9.11 | 0.67 | 39.39 | 3.35 |
| EUC_TP_281 | 16.85 | 5.62 | 3.37 | 0.67 | 33.61 | 3.39 |
| EUC_TP_282 | 5.02 | 5.62 | 0.97 | 0.67 | 43.34 | 3.35 |
| EUC_TP_283 | 25.12 | 5.62 | 1.74 | 0.67 | 37.88 | 3.38 |
| EUC_TP_284 | 18.95 | 5.61 | 7.30 | 0.67 | 33.68 | 3.25 |
| EUC_TP_285 | 10.99 | 5.62 | 2.20 | 0.67 | 27.95 | 3.37 |

| | | | | | | |
|---|---|---|---|---|---|---|
| EUC_TP_286 | 19.51 | 5.62 | 1.56 | 0.67 | 35.23 | 3.39 |
| EUC_TP_287 | 12.20 | 5.61 | 8.11 | 0.67 | 29.43 | 3.36 |
| EUC_TP_288 | 13.19 | 5.62 | 10.62 | 0.67 | 33.94 | 3.38 |
| EUC_TP_289 | 15.76 | 5.62 | 11.38 | 0.67 | 27.29 | 3.39 |
| EUC_TP_290 | 10.86 | 5.62 | 2.85 | 0.67 | 32.02 | 3.35 |
| EUC_TP_291 | 13.76 | 5.62 | 6.34 | 0.67 | 25.32 | 3.36 |
| EUC_TP_292 | 17.58 | 5.62 | 3.92 | 0.67 | 31.88 | 3.34 |
| EUC_TP_293 | 13.97 | 5.61 | 1.63 | 0.67 | 32.99 | 3.34 |
| EUC_TP_294 | 9.89 | 5.61 | 1.52 | 0.67 | 32.89 | 3.36 |
| EUC_TP_295 | 7.34 | 5.62 | 0.30 | 0.67 | 29.07 | 3.24 |
| EUC_TP_296 | 8.25 | 5.61 | 0.98 | 0.67 | 35.60 | 3.35 |
| EUC_TP_297 | 6.20 | 5.62 | 2.53 | 0.67 | 25.04 | 3.39 |
| EUC_TP_298 | 7.00 | 5.61 | 0.99 | 0.67 | 34.55 | 3.25 |
| EUC_TP_299 | 45.08 | 5.61 | 25.16 | 0.67 | 31.73 | 3.23 |
| EUC_TP_300 | 16.67 | 5.62 | 1.50 | 0.67 | 38.92 | 3.43 |
| EUC_TP_301 | 7.87 | 5.62 | 2.19 | 0.67 | 33.65 | 3.42 |
| EUC_TP_302 | 22.02 | 5.62 | 9.30 | 0.67 | 36.24 | 3.37 |
| EUC_TP_303 | 10.01 | 5.62 | 1.27 | 0.67 | 28.46 | 3.36 |
| EUC_TP_304 | 15.00 | 5.61 | 3.61 | 0.67 | 25.04 | 3.35 |
| EUC_TP_305 | 12.97 | 5.62 | 4.81 | 0.67 | 31.32 | 3.42 |
| EUC_TP_306 | 12.34 | 6.48 | 2.40 | 0.78 | 31.43 | 3.30 |
| EUC_TP_307 | -0.37 | 5.62 | 0.02 | 0.67 | 27.76 | 3.34 |
| EUC_TP_308 | 7.53 | 5.62 | 1.33 | 0.67 | 30.15 | 3.34 |
| EUC_TP_310 | 10.84 | 5.61 | 5.93 | 0.67 | 37.71 | 3.36 |
| EUC_TP_311 | 6.04 | 5.61 | 0.46 | 0.67 | 26.74 | 3.35 |
| EUC_TP_312 | 9.00 | 5.62 | 2.33 | 0.67 | 33.82 | 3.24 |
| EUC_TP_313 | 23.39 | 5.62 | 13.68 | 0.67 | 26.27 | 3.33 |
| EUC_TP_315 | 14.36 | 5.62 | 9.12 | 0.67 | 35.03 | 3.26 |
| EUC_TP_316 | 25.25 | 5.62 | 7.85 | 0.67 | 30.80 | 3.34 |
| EUC_TP_317 | 39.44 | 5.62 | 31.39 | 0.67 | 35.38 | 3.24 |
| EUC_TP_318 | 29.86 | 5.61 | 19.11 | 0.67 | 20.85 | 3.35 |
| EUC_TP_319 | 1.47 | 5.62 | 1.53 | 0.67 | 28.39 | 3.24 |
| EUC_TP_320 | 26.39 | 5.61 | 15.76 | 0.67 | 23.91 | 3.34 |
| EUC_TP_321 | 28.30 | 5.62 | 17.52 | 0.67 | 24.46 | 3.35 |
| EUC_TP_322 | 24.42 | 5.61 | 13.10 | 0.67 | 25.50 | 3.35 |
| EUC_TP_324 | 12.89 | 5.61 | 9.19 | 0.67 | 33.18 | 3.24 |
| EUC_TP_325 | 25.11 | 5.62 | 10.30 | 0.67 | 27.44 | 3.36 |
| EUC_TP_326 | 30.95 | 5.62 | 8.67 | 0.67 | 38.00 | 3.35 |
| EUC_TP_327 | 26.76 | 5.62 | 21.83 | 0.67 | 31.01 | 3.34 |
| EUC_TP_328 | 6.29 | 5.62 | 2.29 | 0.67 | 32.28 | 3.23 |
| EUC_TP_329 | 18.61 | 5.62 | 13.88 | 0.67 | 40.36 | 3.34 |
| EUC_TP_330 | 25.94 | 5.61 | 14.96 | 0.67 | 44.42 | 3.24 |
| EUC_TP_331 | 11.06 | 5.61 | 1.45 | 0.67 | 37.18 | 3.34 |
| EUC_TP_332 | 27.49 | 5.62 | 17.46 | 0.67 | 29.99 | 3.35 |
| EUC_TP_333 | 31.85 | 5.62 | 19.50 | 0.67 | 36.68 | 3.36 |
| EUC_TP_334 | 22.13 | 5.62 | 11.37 | 0.67 | 31.10 | 3.35 |
| EUC_TP_335 | 16.63 | 5.62 | 9.48 | 0.67 | 40.38 | 3.34 |

| | | | | | | |
|---|---|---|---|---|---|---|
| EUC_TP_336 | 19.85 | 5.62 | 12.37 | 0.67 | 29.91 | 3.34 |
| EUC_TP_337 | 15.44 | 5.62 | 5.46 | 0.67 | 34.85 | 3.34 |
| EUC_TP_338 | 26.17 | 5.62 | 20.94 | 0.67 | 40.33 | 3.23 |
| EUC_TP_339 | 18.63 | 5.62 | 11.22 | 0.67 | 46.26 | 3.38 |
| EUC_TP_340 | 22.84 | 5.62 | 11.01 | 0.67 | 42.02 | 3.34 |
| EUC_TP_341 | 21.38 | 5.62 | 10.21 | 0.67 | 31.46 | 3.35 |
| EUC_TP_342 | 21.50 | 5.62 | 10.27 | 0.67 | 34.40 | 3.23 |
| EUC_TP_343 | 21.14 | 5.62 | 13.82 | 0.67 | 32.17 | 3.34 |
| EUC_TP_344 | 30.77 | 5.62 | 18.84 | 0.67 | 29.11 | 3.33 |
| EUC_TP_345 | 24.74 | 5.62 | 8.12 | 0.67 | 30.45 | 3.40 |
| EUC_TP_346 | 22.23 | 5.62 | 10.92 | 0.67 | 45.13 | 3.34 |
| EUC_TP_347 | 19.41 | 5.62 | 11.84 | 0.67 | 38.73 | 3.35 |
| EUC_TP_348 | 24.22 | 5.61 | 11.11 | 0.67 | 43.92 | 3.36 |
| EUC_TP_349 | 22.53 | 5.62 | 9.88 | 0.67 | 30.29 | 3.35 |
| EUC_TP_350 | 15.85 | 5.61 | 3.48 | 0.67 | 46.01 | 3.35 |
| EUC_TP_351 | 23.54 | 5.61 | 12.05 | 0.67 | 43.00 | 3.24 |
| EUC_TP_352 | 5.06 | 5.61 | 0.64 | 0.67 | 44.90 | 3.36 |
| EUC_TP_353 | 8.26 | 5.61 | 2.62 | 0.67 | 48.88 | 3.24 |
| EUC_TP_354 | 4.97 | 5.62 | 0.44 | 0.67 | 43.18 | 3.35 |
| EUC_TP_355 | 5.16 | 5.62 | 0.22 | 0.67 | 38.42 | 3.39 |
| EUC_TP_356 | 0.71 | 5.62 | 0.65 | 0.67 | 39.85 | 3.39 |
| EUC_TP_357 | 13.96 | 5.62 | 4.47 | 0.67 | 41.98 | 3.26 |
| EUC_TP_358 | 5.86 | 5.62 | 1.20 | 0.67 | 37.90 | 3.37 |
| EUC_TP_359 | 4.11 | 5.62 | 1.26 | 0.67 | 44.24 | 3.37 |
| EUC_TP_360 | 0.80 | 5.61 | 0.01 | 0.67 | 36.99 | 3.37 |
| EUC_TP_361 | 14.37 | 5.61 | 5.72 | 0.67 | 33.76 | 3.37 |
| EUC_TP_362 | 19.90 | 5.62 | 12.24 | 0.67 | 33.29 | 3.34 |
| EUC_TP_363 | 9.58 | 5.62 | 3.23 | 0.67 | 33.77 | 3.36 |
| EUC_TP_364 | 15.26 | 5.61 | 6.26 | 0.67 | 43.74 | 3.35 |
| EUC_TP_365 | 8.37 | 5.62 | 2.76 | 0.67 | 34.71 | 3.36 |
| EUC_TP_366 | 12.16 | 5.62 | 2.36 | 0.67 | 48.93 | 3.34 |
| EUC_TP_367 | 22.35 | 5.62 | 14.79 | 0.67 | 19.68 | 3.35 |
| EUC_TP_368 | 24.86 | 5.62 | 11.13 | 0.67 | 33.96 | 3.37 |
| EUC_TP_369 | 44.33 | 5.62 | 37.74 | 0.67 | 28.15 | 3.36 |
| EUC_TP_370 | 29.39 | 5.62 | 18.00 | 0.67 | 29.74 | 3.34 |
| EUC_TP_371 | 24.20 | 5.62 | 13.72 | 0.67 | 28.78 | 3.34 |
| EUC_TP_372 | 39.23 | 5.62 | 33.55 | 0.67 | 28.28 | 3.35 |
| EUC_TP_373 | 44.89 | 5.62 | 25.59 | 0.67 | 26.00 | 3.34 |
| EUC_TP_374 | 34.04 | 5.62 | 22.22 | 0.67 | 27.68 | 3.35 |
| EUC_TP_375 | 31.53 | 5.62 | 20.87 | 0.67 | 30.75 | 3.34 |
| EUC_TP_376 | 25.43 | 5.62 | 18.13 | 0.67 | 28.81 | 3.35 |
| EUC_TP_377 | 26.48 | 5.62 | 15.56 | 0.67 | 28.54 | 3.35 |
| EUC_TP_378 | 23.63 | 5.61 | 10.62 | 0.67 | 30.50 | 3.34 |
| EUC_TP_379 | 37.57 | 5.62 | 23.46 | 0.67 | 35.81 | 3.35 |
| EUC_TP_380 | 35.01 | 5.62 | 15.97 | 0.67 | 34.32 | 3.35 |
| EUC_TP_381 | 26.48 | 5.62 | 18.36 | 0.67 | 35.76 | 3.34 |
| EUC_TP_382 | 31.52 | 5.61 | 21.67 | 0.67 | 29.66 | 3.34 |

| | | | | | | |
|---|---|---|---|---|---|---|
| EUC_TP_383 | 10.47 | 5.62 | 3.00 | 0.67 | 34.67 | 3.39 |
| EUC_TP_384 | 31.90 | 5.62 | 16.79 | 0.67 | 32.18 | 3.35 |
| EUC_TP_385 | 23.73 | 5.62 | 16.74 | 0.67 | 34.40 | 3.22 |
| EUC_TP_386 | 38.65 | 5.62 | 30.65 | 0.67 | 28.78 | 3.35 |
| EUC_TP_387 | 39.62 | 5.62 | 28.84 | 0.67 | 31.28 | 3.24 |
| EUC_TP_388 | 31.56 | 5.62 | 17.77 | 0.67 | 34.86 | 3.35 |
| EUC_TP_389 | 17.65 | 5.62 | 8.98 | 0.67 | 27.62 | 3.35 |
| EUC_TP_390 | 45.24 | 5.62 | 28.50 | 0.67 | 29.72 | 3.35 |
| EUC_TP_391 | 18.69 | 5.62 | 8.53 | 0.67 | 28.87 | 3.36 |
| EUC_TP_392 | 12.50 | 5.61 | 5.25 | 0.67 | 34.12 | 3.36 |
| EUC_TP_393 | 16.90 | 5.61 | 5.89 | 0.67 | 44.38 | 3.36 |
| EUC_TP_394 | 20.84 | 5.62 | 14.60 | 0.67 | 37.95 | 3.35 |
| EUC_TP_395 | 9.36 | 5.62 | 4.76 | 0.67 | 34.92 | 3.34 |
| EUC_TP_396 | 16.78 | 5.61 | 3.52 | 0.67 | 34.33 | 3.36 |
| EUC_TP_397 | 22.72 | 5.62 | 12.01 | 0.67 | 47.18 | 3.36 |
| EUC_TP_398 | 42.09 | 5.62 | 28.03 | 0.67 | 41.14 | 3.23 |
| EUC_TP_399 | 14.38 | 5.61 | 6.25 | 0.67 | 37.28 | 3.34 |
| EUC_TP_400 | 8.74 | 5.61 | 0.35 | 0.67 | 28.72 | 3.34 |
| EUC_TP_446 | 10.57 | 5.62 | 1.83 | 0.67 | 33.13 | 3.35 |
| EUC_TP_447 | 31.04 | 5.62 | 6.87 | 0.67 | 32.87 | 3.53 |
| EUC_TP_449 | 16.37 | 5.62 | 10.52 | 0.67 | 29.43 | 3.40 |
| EUC_TP_454 | 8.57 | 5.62 | 1.50 | 0.67 | 26.28 | 3.26 |
| EUC_TP_456 | 6.08 | 5.62 | 0.58 | 0.67 | 24.40 | 3.38 |
| EUC_TP_495 | 68.99 | 5.62 | 59.27 | 0.67 | NA | NA |
| EUC_TP_496 | 71.76 | 5.62 | 58.55 | 0.67 | NA | NA |
| EUC_TP_497 | 67.12 | 5.62 | 53.62 | 0.67 | NA | NA |
| EUC_TP_498 | 68.62 | 5.62 | 51.90 | 0.67 | NA | NA |
| EUC_TP_499 | 79.92 | 5.61 | 73.48 | 0.67 | NA | NA |
| EUC_TP_660 | 28.90 | 5.61 | 19.71 | 0.67 | 44.59 | 3.63 |
| EUC_TP_661 | 7.52 | 5.62 | 0.01 | 0.67 | 43.29 | 3.36 |
| EUC_TP_662 | 22.91 | 5.61 | 16.89 | 0.67 | 44.89 | 3.35 |

[a]square root transformed
[b]back transformed
